# Active sensing during a visual perceptual decision-making task

**DOI:** 10.1101/2025.08.27.672629

**Authors:** Naureen Ghani, Angela Yaxuan Yang, International Brain Laboratory

## Abstract

From the Welsh tidy mouse to the New York City pizza rat, movement reveals rodent intelligence. Here, we show that some head-fixed mice developed an active sensing strategy in a visual perceptual decision-making task. Akin to humans shaking a computer mouse to find the cursor on a screen, some mice wiggled a wheel that controlled the movement of a visual stimulus preferentially during low-contrast trials. When mice wiggled the wheel, the low visual stimulus contrast accuracy increased. Moreover, these wiggles moved the visual stimulus at a temporal frequency (11.5 ± 2.5 Hz) within the range that maximizes contrast sensitivity in rodents. Perturbing the task contingency and visuo-motor coupling reduced wiggle behavior. The performance benefit of wiggle behavior persisted after controlling for arousal state, establishing wiggling as an active sensing strategy rather than an arousal-driven byproduct. Together, these results show that some mice wiggle the wheel to boost the salience of low visual contrast stimuli. This provides evidence for active sensing in head-fixed mouse vision.

**Highlights:**

- Wiggle speed positively correlates with low-contrast visual accuracy across 213 mice
- Wiggles generate stimulus motion at a temporal frequency near maximal contrast sensitivity
- Reversing task contingency or uncoupling the wheel suppresses wiggling
- Longer wiggles are associated with enhanced neural decoding of stimulus identity in midbrain and thalamus

## 1 Introduction

Active sensing is the use of movement to enhance sensory perception and is observed across species and modalities, including chemotaxis in worms,^1^ visual sampling in flies^2,3^, and whisking and sniffing in rats.^4,5^ A common principle of active sensing is moving the stimulus into a region of high acuity.^6^ For example, visual active sensing in humans and monkeys relies on the fovea, which is the high-acuity region of the retina. When humans perform eye fixations, these miniature movements position visual objects onto the fovea and enhance discrimination of fine spatial detail.^7^ Furthermore, micro-saccades modulate the temporal frequency of visual stimuli, which increases visual acuity in humans^8,9^ and monkeys.^10–12^

Because rats and mice lack a fovea, active sensing in rodents has primarily been studied in the somatosensory and olfactory systems rather than vision. Rodents use whisking to locate objects in space^13,14^ and sniffing to sample olfactory information.^15,16^ Active sampling of touch and smell in rodents involves coordinated movements of head bobbing, nose twitching, whisking, and breathing^4,5^. These rhythmic movements are supported by pre-motor oscillator networks in the brainstem.^17–19^

Most rodent studies of active sensing in vision are in freely-moving animals.^20^ During natural visual behaviors, mice coordinate eye, head, and body movements to detect threats, ^21,22^ capture prey,^23^ and judge depth.^24,25^ When mice explore an arena with spots varying in spatial and temporal frequencies, neurons in primary visual cortex (V1) respond most strongly to gaze shifts.^26^ Similar gaze-shift responses are observed in the superficial layers of the superior colliculus (sSC), whereas deeper layers (dSC) instead exhibit motor-related activity aligned to head movements.^27^ This suggests that gaze-shifting head and eye movements follow a dynamic temporal sequence^28^ in neural activity as mice engage in naturalistic active vision.

In contrast, much less is known about active sensing in vision under head-fixed conditions. During visual tasks, head-fixed mice often perform spontaneous, uninstructed movements. Such movements increase the signal-to-noise ratio of V1 responses^29^ and are associated with high arousal levels.^30^ Moreover, measures of arousal and spontaneous movement can predict task engagement^31^ and choice biases.^32^ However, it remains unclear whether such movements can be used by head-fixed rodents for actively sampling visual information.

Here, we propose that head-fixed mice develop an active sensing strategy during a visual perceptual decision-making task. We hypothesize that when visual information is weak, mice deliberately wiggle a steering wheel that controls the visual stimulus position to generate rapid back-and-forth movements. These movements convert a static visual stimulus into a dynamic one. Since the mouse visual system is sensitive to motion,^33^ these mouse-generated wiggles can increase the salience of the visual stimulus.

We test this hypothesis using large-scale data analyses across and within mice. We ask (1) does the low visual stimulus contrast accuracy increase with wiggles? This question is motivated by examples of visual accuracy gains with head-movement jitters^34^ and retinal-image jitters^35,36^ in humans. (2) Are wiggles explained by arousal or change-of-mind? This question is motivated by examples of head-fixed mice fidgeting when aroused^37^ and monkeys reversing movement trajectories with more evidence during decision-making tasks.^38^ (3) Are higher speed wiggles more accurate? This question is motivated by studies that demonstrate improvement in psychophysical performance with higher speed visual stimuli in humans^39^ and monkeys.^40^ Motivated by prior work on active sensing in freely moving rodents, we used continuous and discrete metrics of wiggles to determine whether head-fixed mice use active sensing for vision.

## 2 Results

### 2.1 Mice used a wheel to center a visual stimulus for sugar-water reward

We analyzed data from 213 mice that performed a visual perceptual decision-making task while head-fixed (**Figure 1a**). Mice were trained to use a miniature steering wheel that controlled the movement of a visual stimulus.^1^ When mice moved the wheel for a total of 8.75 mm to center the visual stimulus, they earned sugar-water reward (**Figure 1b**). With a visuo-motor gain of 4 visual degrees per mm of wheel movement, this 8.75 mm corresponded to 35 visual degrees of stimulus movement (8.75 mm ×4 visual degrees/mm = 35 visual degrees).

**Figure 1:**
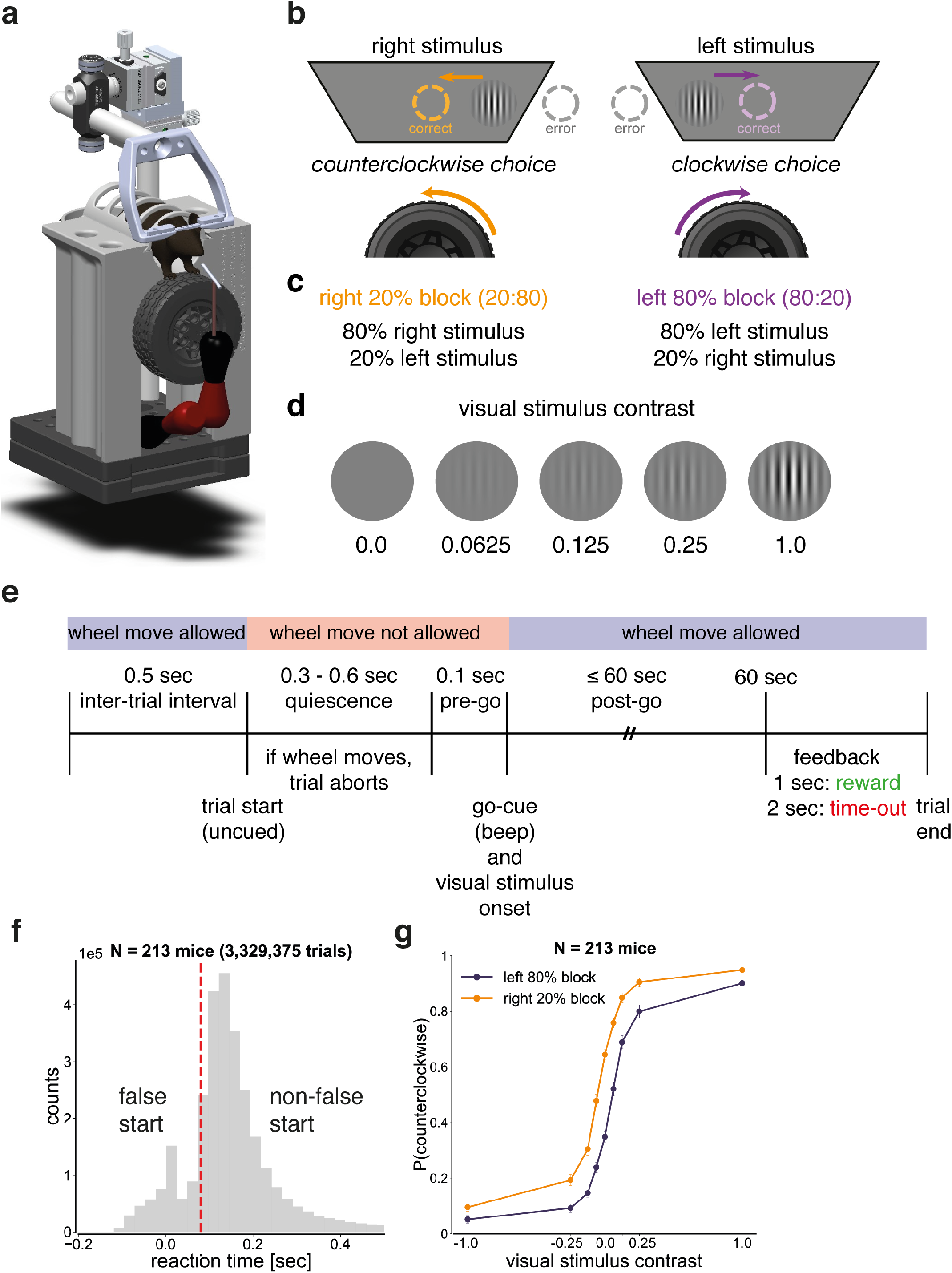
Mice used a wheel to center a visual stimulus for reward. **(a)** Experimental set-up. Head-fixed mice used a miniature steering wheel to report perceptual decisions.^41^ Illustration credit: Florian Rau. **(b)** Task design. Mice turned the wheel counterclockwise for rightward visual stimuli and clockwise for leftward visual stimuli to obtain a sugar-water reward. During error trials, in which the visual stimulus moved off-screen, mice received negative feedback (a 2-second time-out with a gray screen). **(c)** Block structure. In left-biased blocks, 80% of trials contained leftward visual stimuli and 20% contained rightward visual stimuli (80:20). In right-biased blocks, 80% of trials contained rightward visual stimuli and 20% contained leftward visual stimuli (20:80). **(d)** Visual stimuli. Mice were presented with a Gabor patch (orientation: 0°; spatial frequency: 0.10 cycles per degree; Gaussian window: 7 square degrees) at one of five contrast levels (0%, 6.25%, 12.5%, 25%, or 100%). **(e)** Trial structure. At the start of each trial, mice were required to remain still for a quiescence period of 0.4 - 0.7 seconds, which they self-initiated. Upon completion of the quiescence period, a visual stimulus appeared on either the left or right side of the screen (±35° azimuth), accompanied by a 100 msec, 5 kHz sine wave. Mice then had a free-response period of up to 60 seconds to center the visual stimulus for reward. **(f)** Reaction time distribution (mean: 0.69 seconds, median: 0.16 seconds, standard deviation: 2.15 seconds). The red dashed line indicates the threshold for a visual stimulus-triggered response (0.08 seconds). Trials with a reaction time ≤ 0.08 seconds were classified as false starts, whereas all other trials were classified as non-false starts. Reaction time data exceeding 0.5 seconds (13.90% of trials) are not shown. **(g)** Psychometric curves for false start and non-false start trials. In non-false starts trials, the proportion of counterclockwise wheel movements depended on visual contrast (one-way repeated measures ANOVA, left-biased blocks: F = 982.3, p < 0.001, right-biased block: F = 1109.7, p < 0.001) and block type (one-way repeated measures ANOVA, p < 0.05) in non-false starts. Error bars denote the binomial proportion confidence intervals (CI).

We used a shaping protocol to gradually train mice to perform the visual decision-making task at increasingly difficult contrast levels. Mice began with trainingChoiceWorld sessions (left and right stimuli were presented with equal probability)^42^ containing only 100% and 50% contrast trials. When the accuracy reached 70%, we consecutively added lower contrast trials: first adding 25% contrast, then adding 12.5% contrast while replacing 50% with 0% contrast trials and finally adding 6.25% contrast trials.

After learning the visual detection task, mice progressed to biasedChoiceWorld sessions (left and right stimuli were presented with unequal probability). When neural data were acquired during these biasedChoiceWorld sessions, we refer to them as ephysChoiceWorld sessions. Our behavioral dataset consisted of 89% biasedChoiceWorld sessions (2,761 sessions) and 11% ephysChoiceWorld sessions (340 sessions).

The probability of a left visual stimulus was manipulated in strictly alternating biased blocks for sets of 20-100 trials (**Figure 1c**). A session consisted of (1) 90 trials in an unbiased block (50/50) followed by (2) 4-5 left (80/20) and right (20/80) blocks. The independent variable was visual stimulus contrast: 100%, 25%, 12.5%, 6.25%, or 0% contrast (**Figure 1d**).

Each trial began with a self-paced quiescent period during which mice were required to withhold wheel movement for a variable duration between 0.40 and 0.70 seconds. This duration was computed as 0.2 seconds plus a value drawn from a truncated exponential distribution (mean: 0.35 seconds, range: 0.20 - 0.50 seconds) (**Figure 1e**). Early wheel movements during this period reset the quiescence time, rather than being strictly disallowed. Completion of the quiescence period triggered stimulus onset. Following trial outcome, the inter-trial interval was at least 0.5 seconds after correct trials and at least 2 seconds after error trials.

The reaction time distribution across mice (3,329,275 trials) is bimodal, which is how we defined 0.08 seconds as the threshold for a stimulus-triggered response (**Figure 1f**). We plotted reaction times between −0.20 and 0.50 seconds, which represents 86.10% of the data. This implies that the proportion of data not shown in **Figure 1f** is 13.90%, where the reaction time exceeded 0.50 seconds. During stimulus-triggered trials in biased blocks, which followed an initial set of 90 trials in an unbiased block at the start of each session, the proportion of counterclockwise choices depended on block identity (**Figure 1g**). Thus, mouse behavior was biased by blocks.

### 2.2 Wiggles are wheel movements with one or more changes in wheel direction

We defined *k* as the number of wheel directional changes during a trial. We computed *k* from wheel position data interpolated at 1000 Hz, capturing the general motion of the wheel along with superimposed rapid directional changes. Directional changes were identified by counting zero crossings in the smoothed derivative of the position signal (20 Hz low-pass Butterworth filter), which corresponded to points where the wheel velocity (in degrees per second) changed sign.

Non-wiggles were defined as trials with *k* = 0 (**Figure 2a**); these are ballistic movements, characterized by high velocity and acceleration and completed within a short time. Wiggles were defined as trials with *k* ≠ 0 (**Figures 2b-f**). During 6.25% contrast trials, wiggle movements accounted for 45.7% of trials (**Figure 2g**).

**Figure 2:**
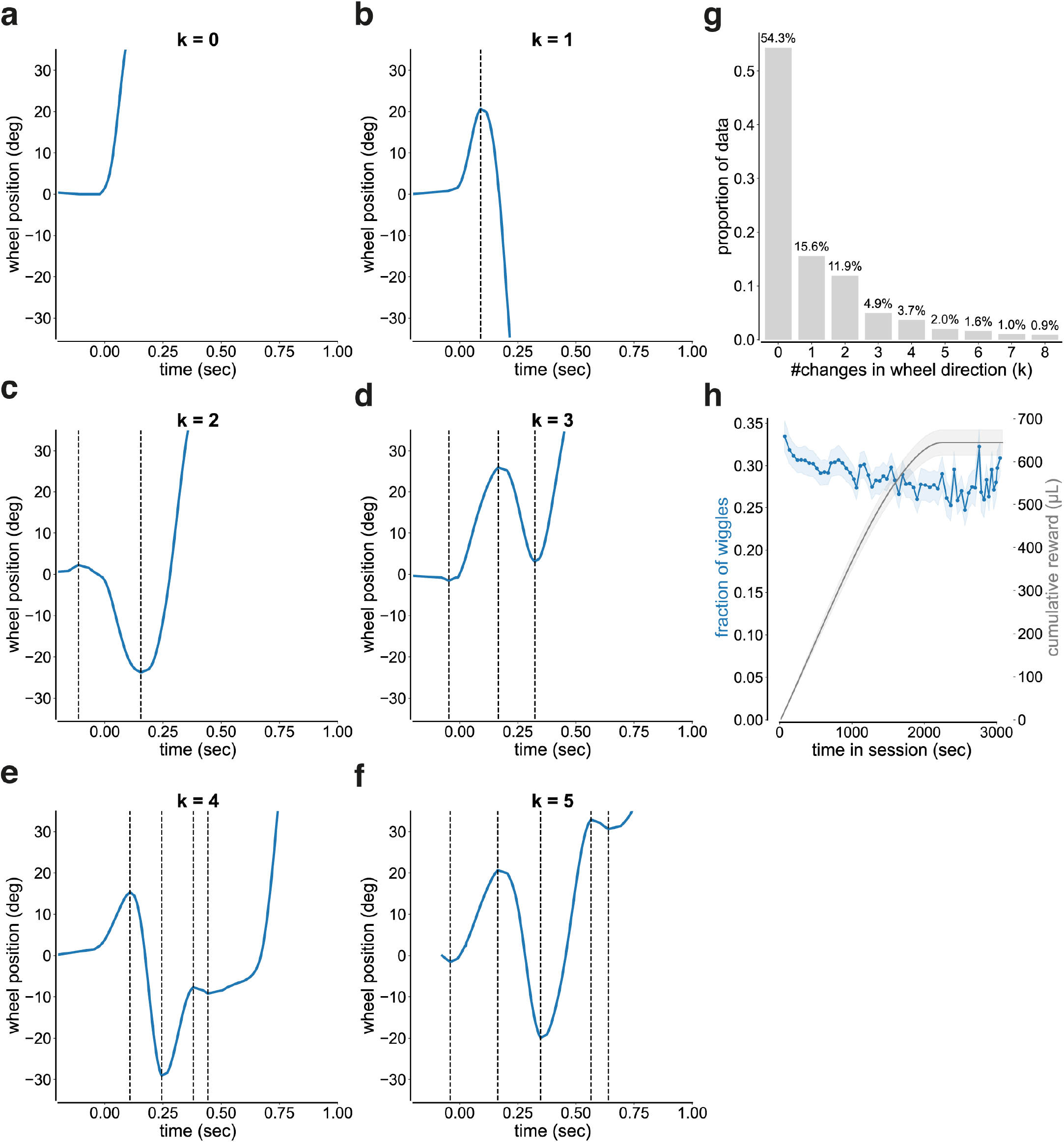
Wiggles are wheel movements with one or more changes in wheel direction. **(a-f)** Single-trial examples of wheel position over time for 6.25% contrast trials with **(a)** *k* = 0, **(b)** *k* = 1, **(e)** *k* = 2, **(d)** *k* = 3, **(e)** *k* = 4, and **(f)** *k* = 5 directional changes. All trials were aligned to motion onset. Only trials with reaction times between 0.08 and 1.20 seconds were included in this and all subsequent analyses. Wiggles were defined as trials with *k* ≠ 0. Non-wiggles were defined as trials with *k* = 0. **(g)** Distribution of the number of directional changes (*k*) for 6.25% contrast trials (mean: 1.75, median: 0.0, standard deviation: 4.50). Trials with more than 8 directional changes (4.13% of trials) are not shown. **(h)** Proportion of wiggle trials over time (500-sec bins; mean ± 95% CI). The orange line indicates the fraction of wiggles, and the gray line indicates the cumulative reward (µL). The fraction of 6.25% contrast wiggle trials decreased over time (linear regression; *β* = −1.17 ×10, R = 0.35, p = 4.8 ×10^7^, N = 61 bins).

We considered three possible interpretations of wiggles: (1) active sensing, (2) change-of-mind, and (3) disengagement.

- **Active sensing hypothesis:** Mice wiggle the wheel to enhance perception because it is easier to detect a moving visual stimulus than a static one. In this view, the jitter serves as a strategy for accumulating sensory evidence by dynamically modulating the visual input.
- **Change-of-mind hypothesis:** Mice wiggle the wheel due to uncertainty about the correct decision. In this view, the jitter represents hesitation between the leftward and rightward choices as the mouse re-evaluates its initial decision.
- **Disengagement hypothesis:** Mice wiggle the wheel because they are disengaged. In this view, the jitter represents idle movements that are not goal-directed.

To address the attentional conditions of wiggle behavior, we quantified the proportion of wiggle trials over time in session. We observed that wiggle behavior decreased as mice became satiated (p = 4.8 ×10^7^; **Figure 2h**). This suggests that wiggle trials are associated with engagement across mice.

### 2.3 Wiggle behavior enhances accuracy at low visual stimulus contrast

#### 2.3.1 The relative contribution of wiggle behavior is greatest during low-contrast trials

To test whether wiggle behavior improves perceptual performance when visual stimulus information is limited, we examined how choice accuracy varied with wiggle speed across visual contrasts. Across mice, faster wiggles were associated with higher choice accuracy at low contrast (+1.9%, d = 1.41; **Figure 3a**), consistent with increased visual salience from rapid stimulus motion. This relationship was reduced for zero-contrast trials (+1.0%, d = 0.76; **Figure 3b**) and near zero for high-contrast trials (+0.0%, d = 0.46; **Figure 3e**), where baseline performance was already close to ceiling. The small gain at zero contrast likely reflects the influence of block prior information from previous trials rather than visual input. A Friedman test confirmed that the strength of the wiggle-accuracy relationship differed across contrast conditions (*χ*^2^ = 49.79, p = 1.54 ×10^11^). Together, these results indicate that the relationship between wiggle behavior and accuracy depends on visual contrast.

**Figure 3:**
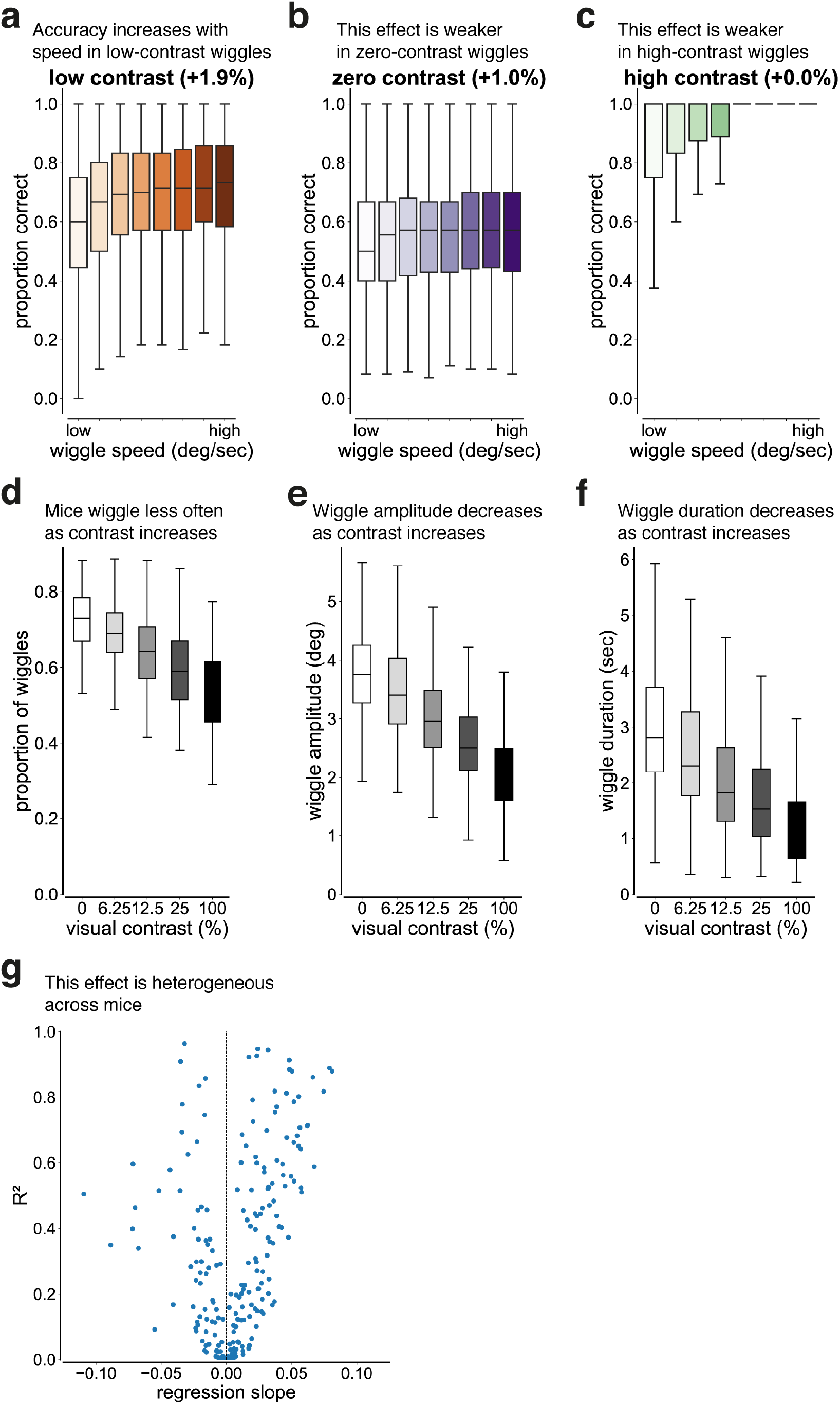
Wiggle behavior enhances accuracy at low visual stimulus contrast. **(a-c)** Distributions of proportion correct as a function of wiggle speed for a given visual contrast. Wiggle speed was defined as the total angular distance between successive directional changes divided by the wiggle duration, which was defined as the time between the first and last extremum. Data were pooled across 213 mice and binned into equal octiles; only sessions with ≥3 trials per bin were included. Percentages in parentheses denote the median change in proportion correct across octiles. Median ± SD number of trials per octile across all mice: 30,083 ± 967 (6.25% contrast), 30,037 ± 990 (0% contrast), 18,628 ± 1,124 (100% contrast). **(a)** Accuracy increased with wiggle speed in 6.25% contrast trials (logistic regression; *β* = 0.063 ± 0.004 SE, p < 0.001, Cohen s d = 1.41, N = 213 mice). **(b)** Accuracy increased slightly with wiggle speed in 0% contrast trials (logistic regression; *β* = 0.026 ± 0.002 SE, p < 0.001, Cohen s d = 0.76, N = 213 mice). **(c)** Accuracy increased modestly with wiggle speed in 100% contrast trials (logistic regression; *β* = 0.18 ± 0.012 SE, p < 0.001, Cohen s d = 0.46, N = 209 mice). Four mice were excluded from this analysis because they did not have enough trials to split into octiles. **(d)** Wiggle occurrence as a function of visual contrast. The proportion of wiggles decreased systematically with increasing visual contrast (linear mixed-effects regression; *β* = −0.16 ± 0.004 SE, p < 0.001, N = 213 mice). **(c)** Wiggle amplitude as a function of visual contrast. The wiggle amplitude decreased systematically with increasing visual contrast (linear mixed-effects regression; *β* = −1.45 ± 0.04 SE, p < 0.001, N = 213 mice). **(f)** Wiggle duration as a function of visual contrast. The wiggle duration decreased systematically with increasing visual contrast (linear mixed-effects regression; *β* = −1.49 ± 0.05 SE, p < 0.001, N = 213 mice). **(g)** Relationship between regression slope and goodness-of-fit in the *k*-analysis. Scatter plot of per-mouse regression slope versus R^2^ for the relationship between proportion correct and the number of directional changes (*k*) during 6.25% contrast trials. Each point represents a mouse, and the black dashed line indicates a slope of zero.

To quantify the relative behavioral benefit of wiggling within individual mice, we computed per-mouse normalized gains in accuracy relative to baseline performance for each contrast condition, where baseline was defined as the accuracy of the lowest wiggle speed bin. Wiggle speed produced the largest relative improvement at low contrast (d = 1.39), a moderate improvement at zero contrast (d = 0.62), and a slight impairment at high contrast (d = −0.36), where baseline performance was near ceiling.

To explore the moderate improvement on zero-contrast trials, we examined the timing of wiggle behavior relative to block switches. We hypothesized that mice remembered the block bias more effectively during faster wiggles. Consistent with this prediction, higher-speed wiggles occurred earlier in the block, with the median number of trials since the most recent block switch decreasing by 3.8% (26.1 ± 3.0 trials since block switch across mice, d = 0.23, p < 0.001). This result suggests that wiggle behavior on zero-contrast trials reflects memory of the block bias rather than sensory input.

Consistent with these observations, post-hoc within-mouse Wilcoxon signed-rank comparisons with Bonferroni correction (*α* = 0.0167) showed that gains at low contrast were significantly larger than those at both zero contrast (W = 3,554, p = 7.06 ×10^17^) and high contrast (W = 6,417, p = 5.85 ×10^7^). These results demonstrate that the magnitude of wiggle-related performance benefit is greatest when visual stimulus information is limited, consistent with wiggle behavior serving as an active sensing strategy to enhance the salience of low-contrast stimuli.

#### 2.3.2 Wiggle behavior cannot be explained by longer decision times

To test whether improved performance on wiggle trials could be explained by longer decision times, we fit a population-level logistic regression model predicting choice accuracy from both the number of changes in wheel direction (*k*) and response duration. Across 506,541 low-contrast trials from 213 mice, the number of changes in wheel direction was a significant positive predictor of accuracy (*β* = 0.0157, z = 6.36, p < 0.001), whereas response duration was not (*β* = 0.0151, z = 1.64, p = 0.10). This indicates that the relationship between wiggle behavior and low-contrast accuracy cannot be explained by longer integration time alone, and instead reflects a specific benefit associated with the wiggle behavior itself.

In addition, we binned the low-contrast data (**Figure S1**) into narrow response duration ranges (**Figure S2b**) and observed that performance remained dependent on the number of wheel directional changes, although these effects did not reach statistical significance within individual duration bins.

#### 2.3.3 Mice with asymmetric wiggle behavior preferentially wiggle on the higher-accuracy side

Next, we quantified how often mice engaged in wiggle behavior as a function of visual contrast. The proportion of wiggle trials decreased systematically with increasing contrast (*β* = −0.16; **Figure 3d**), indicating that mice were more likely to wiggle when visual stimulus information was weak. This contrast dependence supports the interpretation of wiggle behavior as an active sensing strategy.

We identified 56 mice with asymmetrical wiggle behavior across visual contrasts using a Wilcoxon signed-rank test (**Figure S3a**). To test whether these mice selectively wiggle on the side associated with higher low-contrast accuracy, we computed the median fraction of wiggles directed toward the non-preferred side. Across the 56 mice, the median fraction is 0.40 (IQR: 0.36 - 0.44). A one-sample Wilcoxon test confirms that this fraction is significantly less than 0.5 (p < 0.001), indicating that most wiggles are directed toward the more effective side (**Figure S3b**). These results demonstrate that mice predominantly wiggle on the side associated with higher accuracy.

#### 2.3.4 Wiggle behavior is contrast-dependent

We then asked whether the magnitude of wiggle behavior also depended on visual contrast. Wiggle amplitude decreased monotonically with increasing contrast (*β* = −1.45; **Figure 3e**). This means that mice executed larger movements under low-contrast conditions. Because larger-amplitude wheel movements generate stronger visual motion signals, this scaling is expected to enhance stimulus salience. Unlike the foveal visual systems of primates where small-amplitude micro-saccades improve visual perception,^43,45^ the non-foveal mouse visual system benefits from larger stimulus displacements for increasing visual sensitivity.

We next asked whether the duration of wiggle behavior depended on visual contrast. Wiggle duration was defined as the total time spent in rapid, low-amplitude, back-and-forth wheel movements within a trial, excluding periods of stillness or slow drift. If wiggles during low-contrast trials reflect an active sensing strategy, they should be sustained for longer durations, whereas brief wiggles during high-contrast trials would be more consistent with arousal-related jitters. Consistent with this prediction, wiggle duration decreased monotonically with increasing contrast (*β* = −1.49; **Figure 3f**). This indicates that mice sustained wiggle behavior longer when stimulus information was limited.

#### 2.3.5 Wiggle behavior is heterogeneous across mice

To assess whether the benefit of wiggle behavior varied across individual mice, we performed a per-mouse logistic regression analysis between the number of changes in wheel direction (*k*) and choice accuracy during low-contrast trials. Based on a goodness-of-fit criterion (R^2^ ≥0.1), 104 of 213 mice (48.8%) exhibited a positive slope, whereas 52 mice (24.4%) exhibited a negative slope (**Figures 3g, S4a**). Mice with negative slopes that did not benefit from wiggle behavior were more impulsive and had a higher proportion of false start trials (p = 0.001, d = 0.87, Mann-Whitney U test, *α* = 0.0167; **Figures S5, S6**). This suggests that jitters in non-benefiting mice reflected premature or uncontrolled movements rather than deliberative active sensing. Thus, the effect of wiggle behavior on perceptual performance depends on whether jitters are used as an active sensing strategy or as an impulsive motor output.

Across the population, the distribution of slopes was asymmetric and not consistent with random variation: mice with positive slopes had a median slope of +0.029 (IQR: 0.019 - 0.053), whereas mice with negative slopes had a median slope of −0.021 (IQR: −0.034 − −0.017). Moreover, 22% of mice exhibited a strong relationship between *k* and accuracy (R^2^ ≥0.7), indicating a substantial effect in a sizeable subset of the population. These results demonstrate that the impact of wiggle behavior is heterogeneous across mice.

The variance in the R distribution across mice (*σ*^2^ = 0.078) is more than that within mice (*σ*^2^ = 0.0016). Since there is more variation between mice compared to within mice (0.078 > 0.0016), this suggests that there are mice that use wiggles differently from other mice. This aligns well with the generalized linear model-hidden Markov model (GLM-HMM) analysis that distinguishes mice for whom wiggling is helpful versus hurtful for accuracy (**Figure S5**).

### 2.4 Perturbation experiments suggest that wiggle behavior is not driven by change-of-mind

To disambiguate whether wiggle behavior during low-contrast trials reflects an active sensing strategy or a change-of-mind process, we performed three perturbation experiments that altered the coupling between wheel movements and visual feedback. Although these control experiments varied in cohort size, each manipulation preserved the kinematics of wiggle behavior (**Figures S7-9**), while selectively disrupting task contingencies predicted to be irrelevant under a change-of-mind account.

#### 2.4.1 Reversing task contingency suppresses wiggle behavior

First, we trained a separate cohort of 20 mice on a reversed-contingency task in which wheel movements displaced the visual stimulus in the opposite direction. If wiggle behavior reflects active sensing, disrupting the one-to-one mapping between motor output and visual feedback should reduce the performance benefit of wiggling. Although a reversed stimulus-wheel mapping could, in principle, still support wiggling as an active sensing strategy, it runs counter to natural sensorimotor statistics, in which self-generated movements typically produce congruent sensory motion. Such a mismatch is expected to increase sensorimotor prediction error and change how mice use movement to sample sensory information. In contrast, a change-of-mind process predicts that wiggle behavior should persist irrespective of stimulus-action mapping because it reflects internal indecision rather than sensory gain.

In the reversed-contingency task, we observed that wiggle speed was still weakly associated with improved accuracy across contrast conditions (**Figures 4a-e**). However, the magnitude of this relationship was substantially reduced relative to the standard task (GEE logistic regression; task ×wiggle interaction: *β* = −0.088 ± 0.035 SE, p = 0.011; **Figures 3a, 4a**). The absolute increase in accuracy per unit wiggle speed was largest at high contrast (*β* = 0.034, d = 0.75; **Figure 4e**), smaller at low contrast (*β* = 0.026, d = 0.68; **Figure 4a**), and smallest at zero contrast (*β* = 0.015, d = 0.64; **Figure 4b**). Thus, when wheel movements no longer generated veridical stimulus motion, the behavioral benefit of wiggling was strongly reduced. This is consistent with wiggle behavior supporting active sensing rather than reflecting a change-of-mind process.

**Figure 4:**
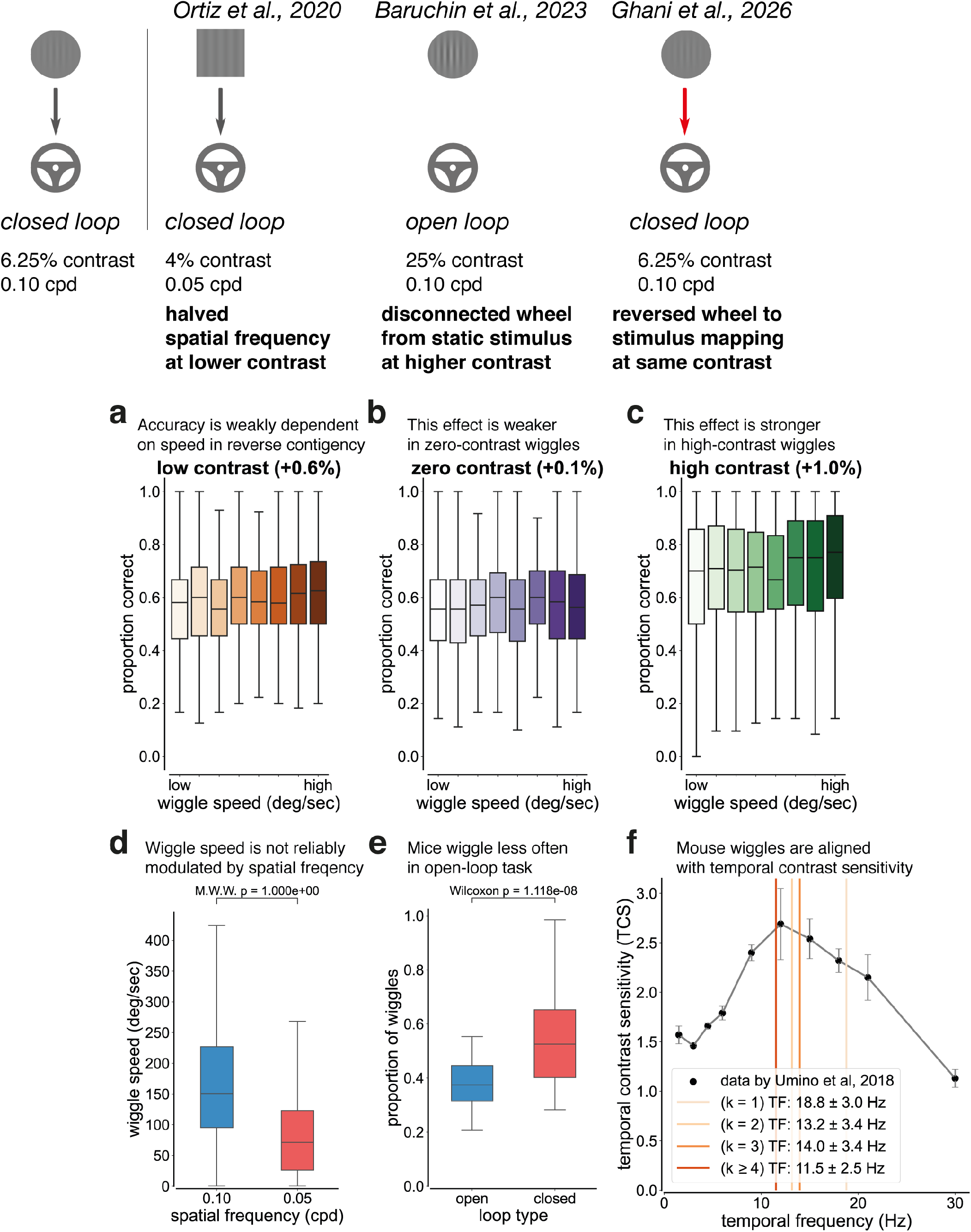
Perturbation experiments suggest that wiggle behavior is not driven by change-of-mind. ***Top:*** Schematic of task variants used as perturbation controls. Arrows indicate that the visual stimulus is yoked to wheel movement (closed-loop), whereas the absence of an arrow indicates that the visual stimulus is uncoupled from wheel movement (open-loop). Gray arrows denote the standard contingency (turning the wheel left moves the visual stimulus left), while red arrows denote a reversed contingency (turning the wheel left moves the visual stimulus right). ***Left-Right:*** Schematics of the original task, spatial frequency control, open-loop control, and reversed-contingency control. **(a)** In the reversed-contingency control, accuracy increased modestly with wiggle speed in 6.25% contrast trials (logistic regression; *β* = 0.026 ± 0.006 SE, p = 0.010, Cohen s d = 0.68, N = 12 mice). **(b)** In the reversed-contingency control, accuracy increased slightly with wiggle speed in 0% contrast trials (logistic regression; *β* = 0.015 ± 0.006 SE, p = 0.012, Cohen s d = 0.64, N = 12 mice). **(c)** In the reversed-contingency control, accuracy increased with wiggle speed in 100% contrast trials (logistic regression; *β* = 0.034 ± 0.012 SE, p = 0.004, Cohen s d = 0.75, N = 16 mice). Four additional mice are included in this analysis because all mice experienced 100% contrast trials. **(d)** Wiggle speed as a function of spatial frequency. The wiggle speed decreased with spatial frequency, but this relationship was not significant (Mann-Whitney-Wilcoxon test; U = 2.0, p > 0.05, N = 1,258 trials, 7 mice, data from Ortiz et al., 2020). Median ± SD values were 150.75 ± 116.66 for 6.25% contrast, 0.10 cpd trials and 71.30 ± 72.20 for 4% contrast, 0.05 cpd trials. **(e)** Wiggle occurrence as a function of loop type. The proportion of wiggles significantly increased from open-loop to closed-loop (Wilcoxon paired samples test, U = 2.0, p = 1.18 ×10^8^, N = 7,515 trials, 4 mice, data from Baruchin et al., 2023). Median ± SD values were 0.37 ± 0.09 for 25% contrast, open-loop trials and 0.53 ± 0.17 for 25% contrast, closed-loop trials. The analysis window was 400 msec for each period, aligned to visual stimulus onset. Each trial consisted of an open-loop period (visual stimulus was uncoupled from the wheel) followed by a closed-loop period (visual stimulus was coupled to the wheel). Mice were rewarded for using the wheel to center the higher contrast Gabor patch (25% versus 100% contrast trials). Wheel movements between stimulus onset and go-cue were ignored. **(f)** Wiggle behavior plotted against temporal contrast sensitivity in the standard-contingency task. Solid lines denote the median temporal frequency, defined as wiggle speed divided by spatial frequency, conditioned on the number of directional changes (*k*). During wiggle trials with *k* ≥4 directional changes, mice moved the wheel at 11.5 ± 2.5 Hz, which is close to the frequency of maximum temporal contrast sensitivity (12 Hz, data by Umino et al., 2018). See **Methods 4.2.4** for details.

#### 2.4.2 Wiggle behavior is not reliably modulated by the spatial frequency of the visual stimulus

Second, we asked whether wiggle speed is modulated by the spatial frequency of the visual stimulus. If wiggle behavior reflects active sensing, then wiggle speed should be inversely proportional to spatial frequency. On the other hand, a change-of-mind process predicts that wiggle speed is not dependent on spatial frequency.

We analyzed data from 7 mice performing a task variant in which the spatial frequency was reduced from 0.10 to 0.05 cycles per degree (cpd).^46^ Median wiggle speed decreased from 150.75 ± 116.66 deg/sec at 0.10 cpd to 71.30 ± 72.20 deg/sec at 0.05 cpd (**Figure 4d**). Although this approximately two-fold reduction in speed when spatial frequency was halved is qualitatively consistent with active sensing, the effect did not reach statistical significance (p > 0.05, N = 1,258 trials, Mann-Whitney-Wilcoxon test). High trial-to-trial variability within individual mice limited our ability to reliably establish spatial frequency modulation of wiggle speed.

#### 2.4.3 Uncoupling the wheel from the visual stimulus suppresses wiggle behavior

Third, we analyzed data from 4 mice performing an open-loop task variant in which wheel movements were disconnected from visual feedback.^47^ When the wheel is uncoupled from the visual stimulus, mice can no longer use wiggling to boost the salience of low-contrast stimuli. If wiggle behavior reflects active sensing, mice should reduce wiggling under open-loop conditions. Consistent with this prediction, we observed that mice wiggled significantly less often during the open-loop period in the lowest contrast condition (25%) (U = 2.0, p = 1.18 ×10^8^, N = 7,515 trials; **Figure 4e**). These results indicate that mouse behavior depends on visuo-motor coupling, supporting the interpretation of wiggling as an active sensing strategy.

Together, these perturbation experiments indicate that wiggle behavior depends on task contingencies and visuo-motor coupling, consistent with an active sensing strategy.

#### 2.4.4 Wiggles generate stimulus motion at a temporal frequency near maximal contrast sensitivity

Lastly, we asked whether wiggle behavior in the standard-contingency task occurs at temporal frequencies aligned with the mouse temporal contrast sensitivity (TCS) function. Mouse TCS was defined as the sensitivity index measured in freely moving mice during a Go/No-Go task.^48^ In that task, the visual stimulus was a full-field Gabor patch (orientation: 0°; spatial frequency: 0.128 cpd; Gaussian window: 7 square degrees; contrast: 50%). Although the visual stimulus in our data is local rather than full-field, the TCS data provides a useful reference for identifying temporal frequencies at which visual motion is most perceptible to mice.

We found that the average temporal frequency of wheel motion during *k* ≥4 wiggle trials (11.5 ± 2.5 Hz) closely matched the frequency at which mouse TCS peaks (12 Hz; TCS = 2.69 ± 0.36 SE; **Figure 4f**). Although *k* ≥4 wiggle trials constitute a small fraction of the data (13.3%; **Figure 2g**), the large overall dataset (25,474 trials from 213 mice) provided sufficient power to estimate the mean temporal frequency generated by wiggle movements (one-sample t-test against 12 Hz: d = 0.196, power = 81.2% at *α* = 0.05). Taken together, these results suggest that mice can perceive the left-to-right visual motion elicited by wiggling. Thus, when mice engage in sustained wiggle behavior, the resulting stimulus motion occurs at temporal frequencies aligned with visual contrast sensitivity.

### 2.5 Arousal modulates the strength of the wiggle-accuracy relationship but does not account for the relationship itself

To determine whether wiggle behavior during low-contrast trials reflects an active sensing strategy rather than differences in arousal, we performed complementary analyses examining the relationship between wiggle behavior, pupil diameter (a widely-used proxy for arousal), and task accuracy.

#### 2.5.1 Wiggle metrics show minimal correlation with pupil diameter

First, we asked whether wiggle metrics were related to pupil diameter across sessions (see **Methods 4.2.l0**). If wiggle behavior is driven by arousal, these relationships should be strong and consistent across metrics. Contrary to this prediction, each of the wiggle metrics (speed, amplitude, number of direction changes, or duration) showed no significant relationship with normalized pupil diameter during either the baseline or post-stimulus period (N = 255 sessions; all p > 0.05; **Figures 5a-d**). These results indicate that wiggle behavior is minimally associated with pupil-linked arousal.

**Figure 5:**
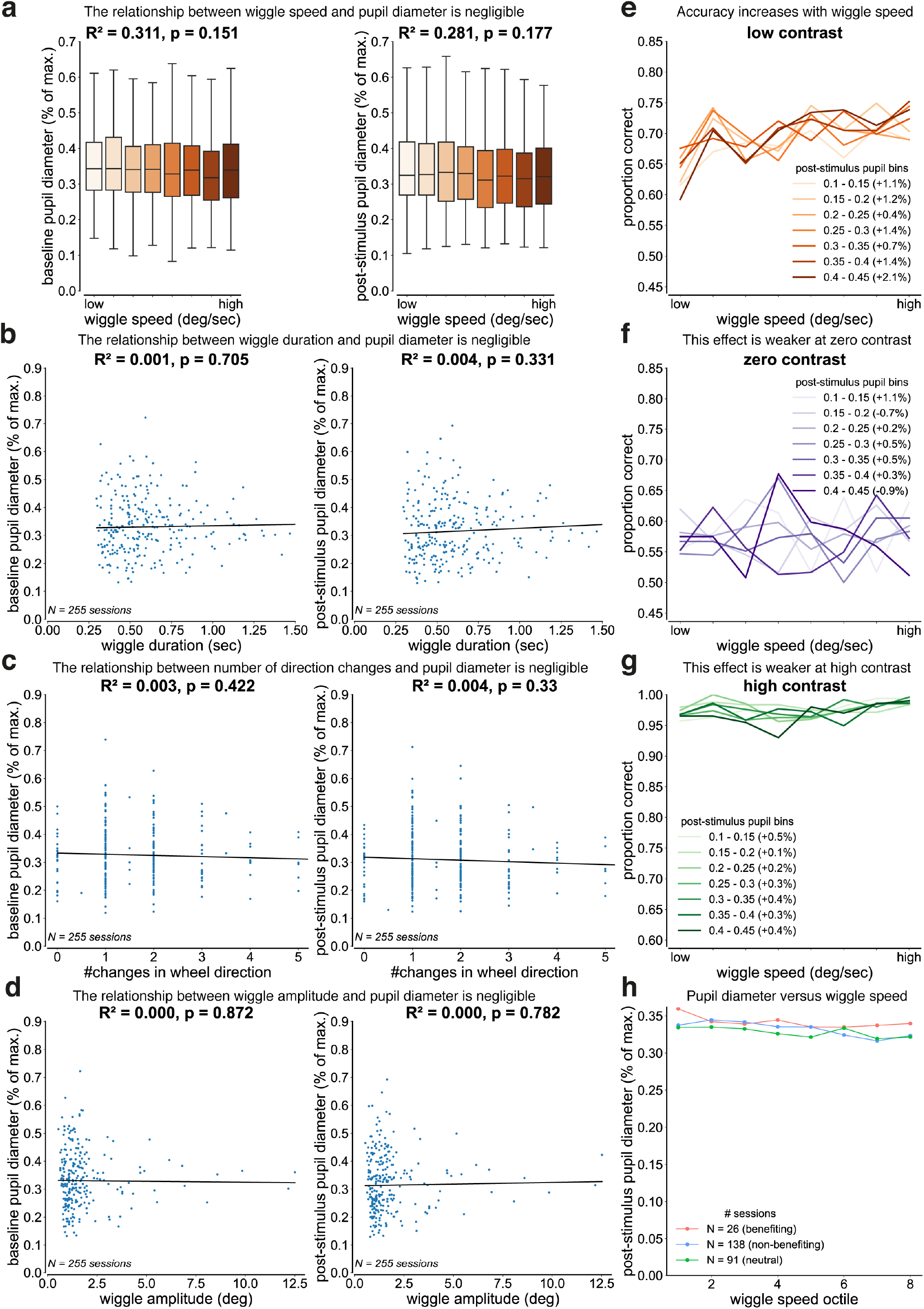
Arousal modulates the strength of the wiggle-accuracy relationship but does not account for the relationship itself. Relationship between wiggle behavior and pupil diameter. Pupil diameter was defined as the vertical distance between the top and bottom pupil coordinates estimated using the Lightning Pose algorithm. For cross-session comparisons, pupil diameter was normalized within each session to the session maximum. Baseline pupil diameter was computed in a 500-ms window before stimulus onset, and post-stimulus pupil diameter in a 500-ms window after stimulus onset. Data were pooled across 255 sessions from 93 mice with manually verified pupil tracking (mean ± SD sessions per mouse: 2.74 ± 1.98). **(a-d)** Session-level relationships between wiggle metrics and normalized pupil diameter during 6.25% contrast trials (N = 255 sessions). For each session, the median of each wiggle metric and the median pupil diameter were calculated across all wiggle trials. Wiggle speed data were binned into equal octiles across sessions, consistent with **Figures 3a and 5e**. R and p-values are from weighted linear regressions, where sessions were weighted by the number of wiggle trials. **(a)** Wiggle speed showed no significant relationship with pupil diameter (baseline: *β* = −0.002, R = 0.311, p = 0.151; post-stimulus: *β* = −0.001, R = 0.281, p = 0.177). **(b)** Wiggle duration showed no significant relationship with pupil diameter (baseline: *β* = 0.010, R = 0.001, p = 0.705; post-stimulus: *β* = 0.027, R = 0.004, p = 0.331). **(c)** Number of direction changes showed no significant relationship with pupil diameter (baseline: *β* = −0.004, R = 0.003, p = 0.422; post-stimulus: *β* = −0.005, R = 0.004, p = 0.33). **(d)** Wiggle amplitude showed no significant relationship with pupil diameter (baseline: *β* = −0.001, R < 0.001, p = 0.872; post-stimulus: *β* = 0.002, R < 0.001, p = 0.782). **(e-g)** Relationship between wiggle speed and accuracy at matched arousal levels (pupil-stratified analysis). Post-stimulus normalized pupil diameter was binned in 0.05 a.u. increments from 0.10 to 0.45. Within each pupil bin, trials were partitioned into octiles of wiggle speed, and accuracy was computed for each octile. Lines show mean accuracy across wiggle speed octiles for each pupil bin. Percentages in legends indicate the mean change in accuracy between successive speed octiles within that bin. R and p-values are from weighted linear regression performed at the trial level within each pupil bin. **(e)** At low contrast (6.25%), accuracy increased with wiggle speed in larger pupil size bins (0.35 - 0.40: R = 0.54, p = 0.03; 0.40 - 0.45: R = 0.60, p = 0.02), with weaker trends at intermediate pupil sizes. **(f)** At zero contrast (0%), no significant relationship between wiggle speed and accuracy was observed in any pupil bin (all R < 0.30, p > 0.05). **(g)** At high contrast (100%), a relationship between wiggle speed and accuracy was present only in the smallest pupil bin (0.10 - 0.15: R = 0.68, p = 0.01) and was absent at larger pupil sizes. **(h)** Relationship between wiggle speed and post-stimulus normalized pupil diameter by session type. Sessions were classified based on the within-session relationship between wiggle behavior (number of wheel direction changes) and low-contrast accuracy using weighted linear regression: benefiting (r > 0.3, N = 26), neutral (−0.3 ≤ r ≤ 0.3, N = 91), or non-benefiting (r < −0.3, N = 138). Trials were binned into wiggle speed octiles within each session. Mixed-effects linear regression with session as a random effect showed no significant relationship between wiggle speed and pupil diameter in benefiting sessions (*β* = −0.002, p = 0.20), but a small significant decrease in neutral (*β* = −0.005, p = 0.02) and non-benefiting sessions (*β* = −0.003, p < 0.001).

#### 2.5.2 Wiggle behavior enhances low-contrast accuracy even when pupil size is held constant

Second, we asked whether the positive relationship between wiggle speed and low-contrast accuracy could be explained by trial-by-trial differences in arousal. If arousal accounts for the wiggle-accuracy relationship, then controlling for pupil size should eliminate this relationship.

##### Pupil-stratified analysis

We stratified trials by post-stimulus pupil diameter into narrow bins and examined the relationship between wiggle speed and accuracy within each bin (see **Methods 4.2.l0**). At low contrast (6.25%), accuracy increased with wiggle speed within individual pupil bins (**Figure 5e**). This effect was strongest in larger pupil size bins (**Table** 1), which suggests that arousal modulates the effectiveness of wiggling as a sensory strategy. Importantly, this effect was specific to low-contrast trials: at zero contrast (0%), no relationship between wiggle speed and accuracy was observed in any pupil bin (**Figure 5f**), and at high contrast (100%), the relationship appeared only in the smallest pupil bin (**Figure 5g**).

**Table 1:**
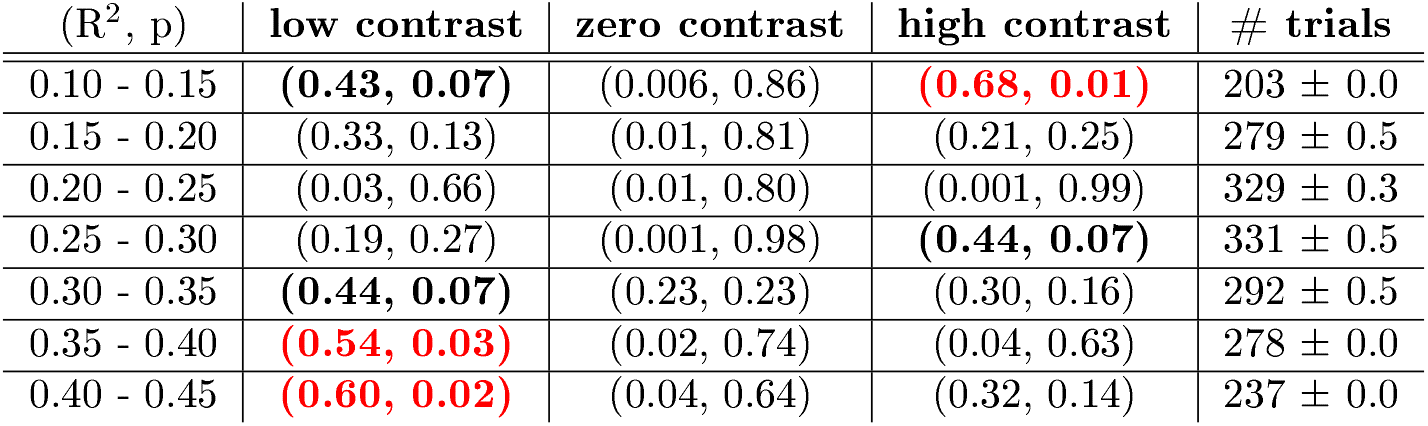
Relationship between wiggle speed and accuraey within pupil bins. R and p-values from weighted linear regressions relating wiggle speed to accuracy within each pupil bin at low (6.25%), zero (0%), and high (100%) contrast. The fourth column reports the mean number of trials per wiggle speed octile across sessions for 6.25% contrast trials (mean ± SD trials). Bold red text indicates R > 0.40 and p < 0.05. Bold black text indicates R > 0.40 and p ≥ 0.05.

**Table 2:** These are caveats in the interpretation of mouse temporal contrast sensitivity.

| visual stimulus properties | The International Brain Laboratory, 2021 | Umino et al, 2018 |
| --- | --- | --- |
| size | local | full field |
| contrast | 6.25% | 50% |
| spatial frequency | 0.1 cycles per degree | 0.128 cycles per degree |
| brightness | 1.47 log cd m<sup>-2</sup> | 1.80 log cd m<sup>-2</sup> |

##### Linear mixed-effects analysis

We used a linear mixed-effects model with both wiggle speed and pupil diameter as fixed effects and mouse identity as a random intercept (see **Methods 4.2.l0**). Wiggle speed significantly predicted accuracy when pupil size was held constant (*β* < 0.001, p < 0.001), while pupil diameter did not predict accuracy when wiggle speed was held constant (*β* = −0.063, p = 0.14) across 93 mice. This confirms that wiggling predicts performance independently of pupil-indexed arousal at the population level.

Together, these results demonstrate that the accuracy benefit associated with wiggle behavior persists even when pupil-linked arousal is controlled for and is selectively expressed under low-contrast conditions, where sensory information is weak. The observation that this effect scales with pupil size suggests that arousal modulates the strength of the wiggle-accuracy relationship but does not account for the relationship itself.

#### 2.5.3 Benefiting and non-benefiting sessions show similar pupil size distributions, and wiggle speed does not increase with pupil diameter in benefiting sessions

Third, we asked whether individual differences in the behavioral benefit of wiggling could be explained by differences in arousal. In **Figure 5a**, we found that wiggle speed showed minimal correlation with pupil diameter when analyzed across all sessions. Here, we examine whether this relationship differs when stratified by session type.

We classified sessions as benefiting, neutral, or non-benefiting based on the within-session relationship between wiggle behavior and low-contrast accuracy (see **Methods 4.2.l0**). While mice tend to exhibit consistent session types over days in training, we performed our analysis at the session level because pupil tracking data were available for only a small subset of sessions per mouse (median: 2 sessions with pupil data per mouse across 93 mice, compared to a median of 19 sessions per mouse across 213 mice in the behavioral analysis). This session-level approach maximizes the statistical power available and tests whether arousal differences explain the heterogeneity in wiggle-related performance benefits.

If arousal explains these differences, we expect that: (1) benefiting sessions should operate at systematically higher arousal levels than non-benefiting sessions, and (2) benefiting sessions should show a positive relationship between wiggle speed and pupil diameter (engagement), while non-benefiting sessions should show a negative relationship (disengagement).

##### Pupil diameter distributions across session types

We compared pupil diameter distributions during low-contrast trials (6.25%) between benefiting sessions (N = 26), neutral sessions (N = 91), and non-benefiting sessions (N = 138) (**Figure S10a**). The three groups showed similar pupil diameter distributions (Kruskal-Wallis test, p = 0.75), with no significant differences in median pupil size (benefiting: 0.34 ± 0.09 a.u.; neutral: 0.29 ± 0.09 a.u.; non-benefiting: 0.31 ± 0.11 a.u.; one-way ANOVA, F(2, 252) = 0.20, p = 0.82). These results indicate that benefiting sessions do not operate at systematically higher arousal levels than non-benefiting sessions.

##### Wiggle speed versus pupil diameter by session type

We binned wiggle speed into octiles (as in **Figures 5e-g**) and examined the relationship between wiggle speed and post-stimulus pupil diameter using mixed-effects linear regression with session as a random effect (see **Methods 4.2.l0**; **Figure 5h**). In benefiting sessions, pupil diameter showed no significant relationship with wiggle speed (*β* = −0.002, p = 0.20). In contrast, neutral and non-benefiting sessions exhibited a small but significant decrease in pupil diameter with increasing wiggle speed (neutral: *β* = −0.005, p = 0.02; non-benefiting: *β* = −0.003, p < 0.001). These results are inconsistent with an arousal-based explanation for benefiting sessions, which predicts that higher-speed wiggles should be associated with higher arousal levels.

Together, these results indicate that wiggle behavior in benefiting sessions enhances performance independent of arousal state, supporting an active sensing strategy rather than an arousal-driven mechanism. The negative relationship observed in neutral and non-benefiting sessions may reflect disengagement during lower arousal states.

### 2.6 Wiggle duration enhances stimulus side decoding accuracy in midbrain and thalamic regions

To test whether wiggle behavior improves the visibility of low-contrast stimuli in neural data, we applied a logistic regression decoder analysis to single-trial, single-neuron electrophysiology data. We hypothesized that stimulus side decoding would be more accurate during wiggle trials than non-wiggle trials. If wiggle behavior reflects active sensing, then mice gain sensory information with each directional change in a wheel movement. We expected individual wiggles to be associated with visual gain and enhanced stimulus side decoding accuracy.

However, one shortcoming of this decoder-based analysis is that the input needs to be the same length from both conditions, and a non-wiggle trial is much faster than a wiggle trial (duration of about 0.2 seconds versus 1 second). The ideal control is long, non-wiggle trials that do not come from losing focus, motivation, or attention. However, such trials do not exist.

We instead asked if the visual stimulus side decoding accuracy increases over the duration of a wiggle trial. We compared decoding accuracy between short (0.2 seconds) and long (1 second) wiggle durations across multiple brain areas.

We show that this is indeed the case in midbrain and thalamic regions. In the midbrain, both the superior colliculus (SC) and anterior pretectal nucleus (APN) showed significant improvements in decoding accuracy with longer wiggles (SC: p < 0.001, +3% accuracy; APN: p < 0.001, +5% accuracy; **Figure 6a**). Similarly, the lateral posterior nucleus of the thalamus (LP) exhibited a significant decoding enhancement (p = 0.001, +3% accuracy; **Figure 6b**). It is important to note that decoding accuracy did not improve with longer wiggles in the primary visual cortex (V1: p = 0.382, −0.7% accuracy; **Figure S11**), but it did in the antero-medial primary visual area (VISa: p = 0.035, +2.2% accuracy; **Figure S11)**.

**Figure 6:**
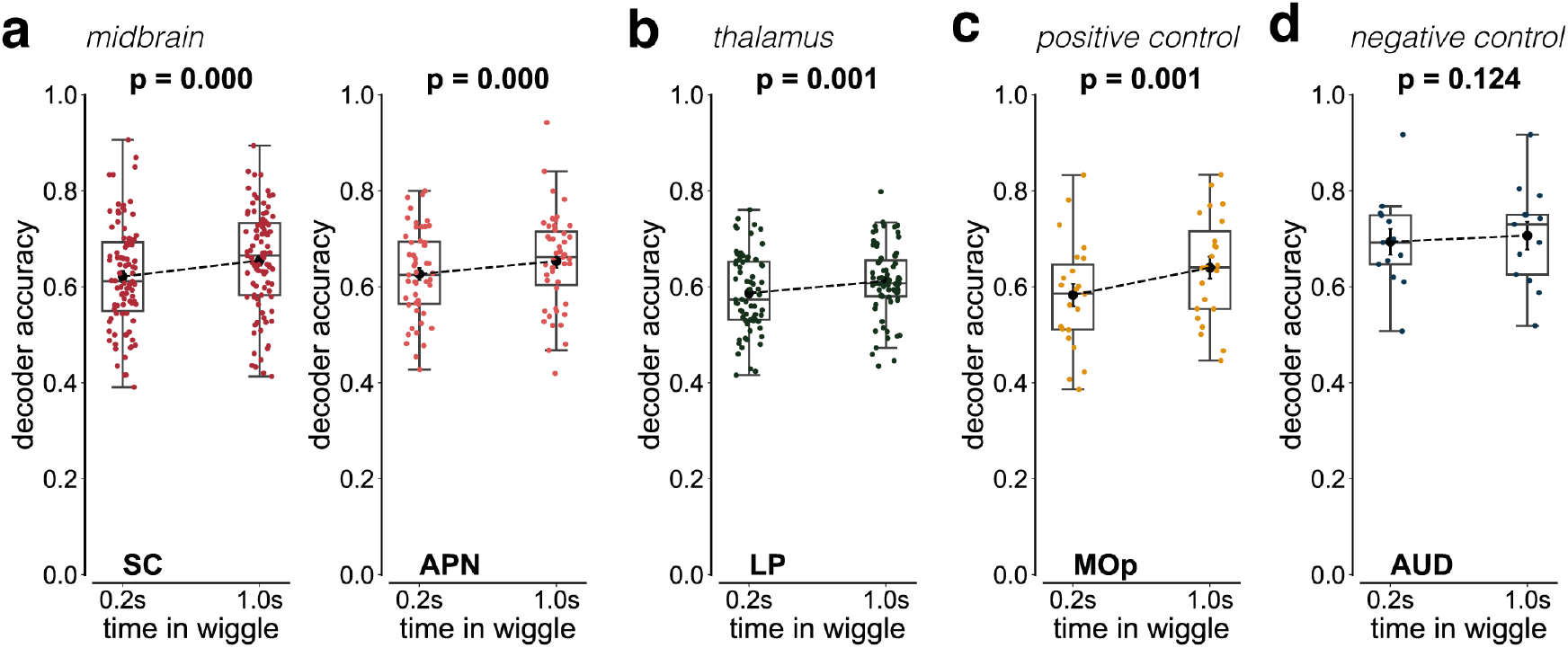
Wiggle duration enhances stimulus side decoding accuracy in midbrain and thalamic regions. A logistic regression decoder was trained to predict visual stimulus side (left versus right) from post-stimulus onset neural activity as a function of wiggle duration (short: 0.2 seconds; long: 1 second). Decoder input consisted of single-trial, single-neuron activity from a single brain region, including only trials with a visual stimulus contrast between 0% and 100%. Decoder output was stimulus side classification accuracy. Only sessions with ≥20 wiggle trials were included. Neural activity was binned in 20-msec time windows, and decoding was repeated 100 times using stratified train-test splits. Brain region abbreviations and color conventions follow the Allen Institute Brain Atlas (SC: superior colliculus; APN: anterior pretectal nucleus; LP: lateral posterior nucleus of the thalamus; MOp: primary motor cortex; AUD: auditory cortex). All statistical comparisons were performed using Wilcoxon signed-rank tests (*α* = 0.05). **(a)** Midbrain regions showed significant improvements in decoding accuracy for long versus short wiggle trials: SC (p < 0.001, N = 91 sessions, median improvement: 3%) and APN (p < 0.001, N = 51 sessions, median improvement: 5%). **(b)** The thalamic LP also showed significant improvement in decoding accuracy (p = 0.001, N = 72 sessions, median improvement: 3%). **(c)** The positive control, MOp, showed a significant improvement in decoding accuracy (p = 0.001, N = 23 sessions, median improvement: 2%). **(d)** The negative control, AUD, showed no significant change in decoding accuracy (p = 0.382, N = 25 sessions, median improvement: 1%).

Positive control analyses in the primary motor cortex (MOp) confirmed expected improvements (p = 0.001, +2% accuracy; **Figure 6e**), whereas the negative control in auditory cortex (AUD) showed no significant change (p = 0.124, +1% accuracy; **Figure 6d**). These results demonstrate that extended wiggle duration selectively improves neural representations of stimulus side in subcortical regions, which is consistent with the hypothesis that longer wiggles provide more visual stimulus information than shorter wiggles.

Together, these results indicate that stimulus side information accumulates over the duration of a wiggle in specific subcortical visual regions (SC, LP) and higher-order visual areas (VISa). This suggests that wiggle behavior increases the visual information content available in those brain regions.

This analysis does not test whether *individual wiggle movements* instantaneously improve stimulus encoding, nor does it disambiguate sensory information gains from longer evidence integration times. Rather, it asks a narrower and more conservative question: whether specific brain regions reflect differences in visual stimulus encoding as a function of wiggle behavior. Indeed, we observed enhanced stimulus side decoding accuracy in midbrain and thalamic regions, but not in auditory cortex. This supports the interpretation that extended wiggle duration preferentially enhances stimulus-related representations in neural circuits relevant for visual processing.

### 2.7 Neuronal firing increases during wiggle trials across multiple brain regions

To relate wiggle behavior to underlying neural activity, we quantified the proportion of neurons with elevated firing rates during wiggle trials compared to non-wiggle trials across 37 brain areas. During limited visual contrast trials, we observed that the top three brain regions modulated by wiggle behavior were: the anteromedial visual area (VISam: 16.67% ± 5.17% SE), superior colliculus (SC: 10.93% ± 2.36% SE), and primary visual cortex (V1: 8.37% ± 2.56% SE) (**Figures 7a-b**). These three brain regions are important for mouse visual processing, which suggests that wiggle behavior engages mouse visual circuits.

**Figure 7:**
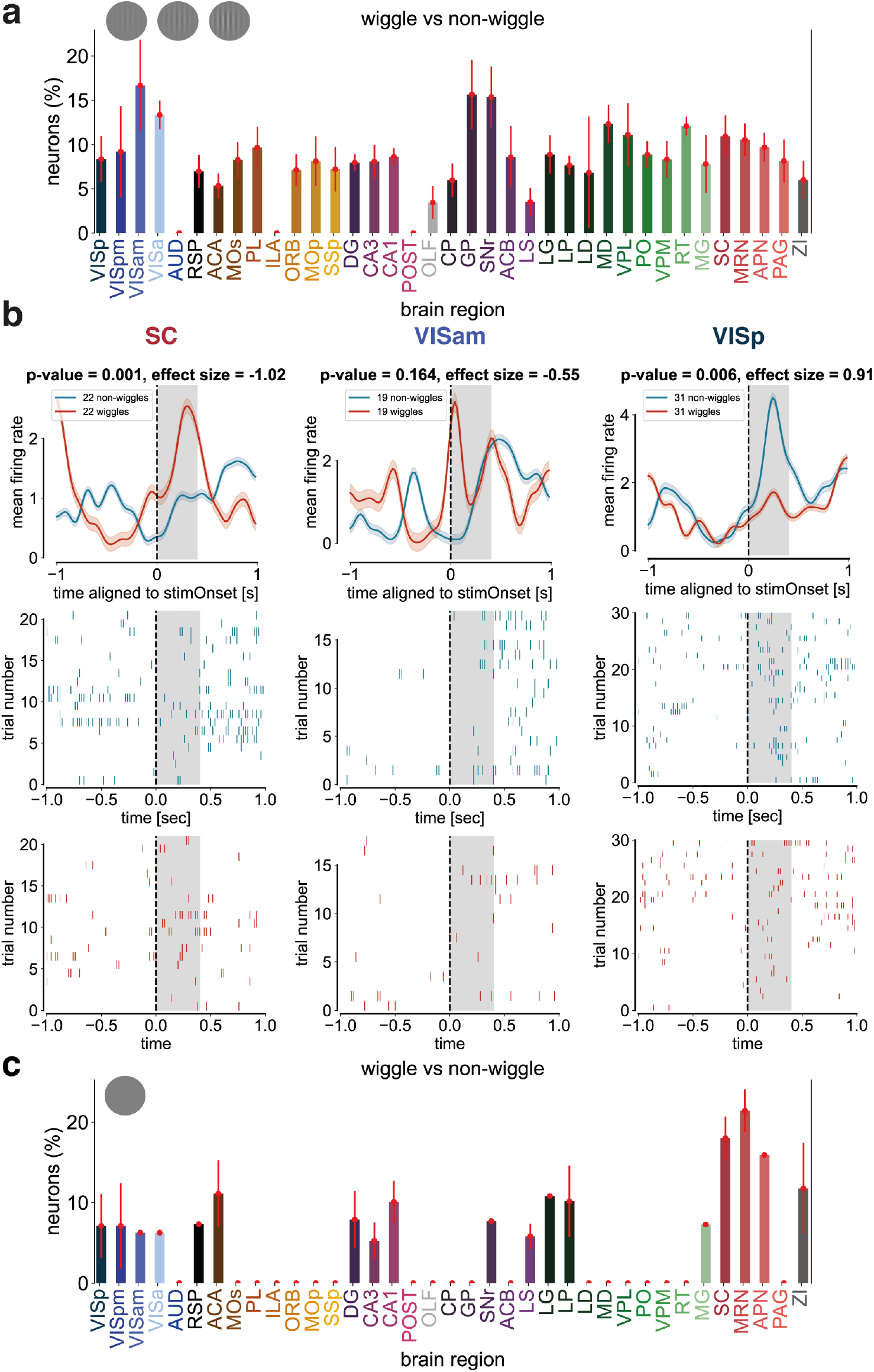
Neuronal firing increases during wiggle trials across multiple brain regions. **(a)** Distribution across 37 brain regions of the fraction of neurons (%) showing a significantly higher normalized mean firing rate during wiggle trials compared to non-wiggle trials in 400-msec window following visual stimulus onset. Only trials with a visual stimulus contrast between 0% and 100% and counterclockwise choice were included. Significance was defined as p ≤ 0.05 (Wilcoxon signed-rank test) and Cohen’s d ≥0.8. Spikes were counted in 20-msec bins, smoothed with a Gaussian kernel (*σ* = 3 bins), and normalized by dividing by baseline firing rate (−0.5 to 0 s relative to stimulus onset) + 1 spike/s. For each wiggle trial, the closest non-wiggle trial in session time was selected. The highest proportion of responsive neurons was observed in the anteromedial visual area (VISam; 16.67%), and the lowest in the olfactory area (OLF; 3.45%). Brain region abbreviations and color conventions follow the Allen Institute Brain Atlas. Error bars denote the standard error of the mean (SEM). **(b)** Example single-neuron responses aligned to visual stimulus onset for the superior colliculus (SC), anteromedial primary visual area (VISam), and primary visual cortex (VISp). **(c)** Negative control showing single-neuron responses during 0% contrast trials.

As a negative control, we performed an identical analysis in the absence of a visual stimulus. During zero-contrast trials, the proportion of wiggle-modulated neurons was reduced in the anteromedial visual area (VISam: 6.25% ± 0% SE) and primary visual cortex (V1: 7.11% ± 3.93% SE), but not the superior colliculus (SC: 18.01% ± 2.66% SE) (**Figure 7e**). This dissociation suggests that visual cortical areas and the superior colliculus play different roles in wiggle behavior.

Taken together with the decoder analysis that showed that stimulus side information increases over the duration of a wiggle in midbrain and thalamic brain regions (**Figure 6**), these firing rate analyses support an interpretation of wiggle behavior as an active sensing strategy. Both analyses indicate that wiggle behavior engages visual brain areas rather than frontal areas, which would suggest a change-of-mind process. If wiggle behavior reflects an active sensing strategy, then extended wheel movements increase the availability of stimulus information in mouse visual circuits.

## 3 Discussion

Our first claim is that a subset of mice uses wheel wiggling as an active sensing strategy to improve visual stimulus detection. This claim is supported by the positive relationship between low-contrast accuracy and wiggle speed across 213 mice, which has a regression line slope that is significantly greater than zero (**Figure 3a**). The large number of trials and animals in our dataset lends confidence to the robustness of the effect. Further support comes from the contrast-dependence of the wiggle behavior itself (**Figures 3d-f**). The systematic increase in the proportion of wiggle trials (**Figure 3d**), wiggle amplitude (**Figure 3e**), and wiggle duration (**Figure 3f**) with decreasing contrast shows that wiggle behavior is dependent on the strength of visual information. Finally, longer wiggle durations are associated with improved decoding of stimulus side in visual brain regions, including the superior colliculus and anteromedial visual area (**Figures 6, 7**). Together, these results support the interpretation that wiggle behavior serves to improve visual stimulus detectability in some mice.

Our second claim is that mouse wheel wiggling cannot be explained by arousal or change-of-mind. To account for arousal, we performed analyses examining the relationship between wiggle behavior and pupil diameter. Wiggle metrics showed minimal correlation with pupil diameter (**Figures 5a-d**), and the positive relationship between wiggle speed and low-contrast accuracy persisted when controlling for arousal through both stratified analysis and mixed-effects regression (**Figures 5e-g, Table 1**). In sessions where wiggling enhances performance, pupil diameter showed no substantial relationship with wiggle speed. In addition, mean pupil diameter was comparable between wiggle and non-wiggle trials (**Figure 3f**) and the median cross-correlation coe cients between pupil data and wheel data were negligible (**Figure S12b**). Notably, the wiggle-accuracy relationship scaled with pupil size: the benefit of wiggling was strongest at higher arousal levels (**Figure 5e, Table 1**). This suggests that arousal modulates the effectiveness of wiggling as a sensory strategy without accounting for the relationship itself, which is consistent with the view that arousal sets a gain on an underlying active sensing mechanism rather than driving it. Together, these analyses demonstrate that wiggle behavior is inconsistent with an arousal-based explanation.

To account for change-of-mind, we reversed the task contingency and observed a decrease in the correlation between low visual stimulus contrast accuracy and wiggle speed across 12 mice (**Figure 4a**). Consistent with this, uncoupling the wheel from the visual stimulus also suppresses wiggling in 4 mice (**Figure 4e**). These perturbation experiments demonstrate that wiggle behavior cannot be explained by change-of-mind.

Our third claim is that mice wiggle the wheel at a natural temporal frequency for the visual system. Active sensing movements are often rapid compared to the speed of sensory transduction itself. Mouse wheel wiggling generates a median visual temporal frequency of 11.5 ± 2.5 Hz during *k* ≥4 directional changes (**Figure 4f**), which is much higher than the 1 - 2 Hz preferred temporal frequency of mouse V1 neurons.^46,47^. However, this visual temporal frequency aligns well with the maximum temporal contrast sensitivity ^45^ of 12 Hz (**Figure 4f**). This finding should be interpreted considering the difference between full-field versus localized visual stimulus.

We show that a subset of mice wiggles the wheel to boost the *salience* of low visual contrast stimuli (**Figure S13**). The salience of an object refers to its color, motion, shape, and brightness. Since the visual stimulus is always oriented vertically during the task, this means that when a mouse jitters the wheel left-to-right, it switches between black and white stripes as the low-contrast Gabor patch moves horizontally on the screen. By wiggling the wheel, head-fixed mice can build optic flow when stimulus information is limited during a visual perceptual decision-making task.

Our work highlights the role of active sensing in head-fixed mouse vision. While ethological studies of naturalistic mouse vision probe at the neural correlates of active eye and head movements,^27,48,49^ it is much harder to do a cell-type specific interrogation of neural circuits in a freely-moving animal compared to a head-fixed animal. Recent advances in virtual reality technology combined with head-fixed recordings offer a promising alternative to measurements in freely moving animals.^50,51^ We demonstrate that head-fixed mice physically wiggle a wheel during low visual contrast stimuli trials in the same spirit of humans performing fixational eye movements during high spatial frequency stimuli trials,^7^ which are both active sensing strategies that serve to increase visual information.

### 3.1 Proposed perturbation experiments

In addition to reversing the task contingency and uncoupling the wheel, there are additional perturbation experiments that probe at different aspects of wiggle behavior.

- **Temporal frequency manipulation:** With a moving stimulus, we hypothesize that mouse wiggle behavior is suppressed.
- **Visuo-motor gain manipulation:** We hypothesize that the amplitude of mouse wiggles scales inversely with the visuo-motor gain. For example, doubling the gain (1 wheel degree ≈ 4 visual degrees) would halve the required wiggle amplitude for the same visual displacement. Because movement velocity depends on amplitude, the perceptual effectiveness of wiggles is constrained by the mouse contrast sensitivity function: movements that are too fast or too slow reduce stimulus visibility.
- **Orientation manipulation:** With a horizontally oriented (90°) stimulus, we hypothesize that mouse wiggle behavior is suppressed. Under this manipulation, we further hypothesize that a subset of mice will adopt an alternative active sensing strategy based on detecting the stimulus edge against the background, thereby partially compensating for the reduced benefit of wiggling.

### 3.2 Limitations of the study

#### 3.2.1 The task was designed to study priors, not active sensing

First, most of the data were acquired during biased blocks. Since mice could exploit the block structure of the task rather than rely solely on the visual stimulus for decisions, this reduces their dependence on wiggles for perceptual enhancement. Second, the spatial frequency of the visual stimulus was above the perceptible threshold. This meant that mice could perceive low visual contrast stimuli without wiggles. Redesigning the task to present more difficult visual stimuli at the perceptual threshold would likely reveal substantially larger benefits from wiggles. Third, the heterogeneous nature of wiggle responses across individual mice means that population averages obscure both mice that benefit from wiggles and those that do not, diluting the overall effect size and masking the biological significance of this behavior (**Figures 3g, S13e**). There are 110 mice with a monotonically increasing correlation, 24 mice with a monotonically decreasing correlation, and 69 mice with a non-monotonically increasing correlation (**Figure S6**).

#### 3.2.2 The spatial frequency perturbation experiment was conducted in 4% contrast stimuli with a rectangular envelope

We show that wiggle behavior is not reliably modulated by the spatial frequency of the visual stimulus (**Figure 4d**). However, these trials used 4% contrast stimuli with a rectangular envelope, whereas the main task used 6.25% contrast stimuli with a circular envelope. A stronger control would be to have mice perform a spatial frequency variant under matched visual stimulus conditions. The hypothesis is that wiggle speed is inversely proportional to spatial frequency.

#### 3.2.3 The open-loop perturbation experiment was conducted in 25% contrast trials

We show that the proportion of wiggle trials decreases when the wheel is uncoupled from the stimulus in 25% contrast trials during a visual discrimination task (**Figure 4e**). A stronger control would be to have mice trained on the original visual detection task with 6.25% contrast trials perform an open-loop variant for consecutive sessions. The hypothesis is that the active sensing strategy would become extinguished over time.

#### 3.2.4 The neural data were not retinotopically aligned to the visual stimulus

For cortical neurons in the retinotopic location of the visual stimulus, a moving visual stimulus would move from one neuron s receptive field to a second neuron s receptive field. This means that two neurons are excited at different times by the dynamic visual stimulus. Thus, a neural circuit mechanism of mouse wiggles likely involves sequentially activating neurons with retinotopically-aligned receptive fields to the visual stimulus. Since the neural data were acquired with a grid-like approach, this sequential excitation cannot be resolved.

Despite this limitation, we observed correlates of wiggles in visual brain areas. In a first-pass analysis, we show that decoding visual stimulus identity from neural activity improves with longer wiggles in midbrain and thalamic regions, including the superior colliculus (**Figures 6, 7**). This result is consistent with prior work showing that the superior colliculus encodes visual representations of active eye movements in head-fixed mice.^52^

In addition to the superior colliculus, a study by Millman and colleagues shows that vasoactive intestinal peptide (VIP) interneurons in mouse V1 were sensitive to low-contrast nasal-to-temporal motion on the retina.^53^ Although this differs from how mouse wheel wiggles move the visual stimulus (left-to-right versus front-to-back), it suggests that VIP interneurons contribute to processing low-contrast retinal motion.

More broadly, these findings in mice align with established neural mechanisms of active vision in non-human primates. During active vision in monkeys, spatiotemporally precise retinal input ^54,55^ is transformed into attention-guided eye movements^56^ through the dorsal cortical pathways to the superior colliculus.^57,58^ While eye movements in mice differ from those of monkeys,^59–61^ we show that a subset of head-fixed mice physically manipulate a wheel to achieve a comparable functional effect. Wheel wiggling therefore provides a behavioral paradigm for elucidating the neural mechanisms of active sensing in head-fixed mouse vision, one of the most well-studied and genetically tractable systems in neuroscience.

## 4 Methods

### 4.1 Experimental model and subject details

All procedures and experiments were carried out in accordance with the local laws and following approval by the relevant institutions: the Animal Welfare Ethical Review Body at University College London; the Institutional Animal Care and Use Committees of Cold Spring Harbor Laboratory, Princeton University, University of Washington, University of California at Berkeley and University of California at Los Angeles; the University Animal Welfare Committee of New York University; and the Portuguese Veterinary General Board.

#### 4.1.1 Mice

A total of 213 adult mice (C57BL6/J, 163 male, 50 female, age: 29.92 ± 13.47 weeks) were used for analysis. Mice were kept in a 12-hour light-dark cycle and performed the task in the dark period (from 7:00 to 19:00). Details are available here: (Appendix 1 of The International Brain Laboratory, 2021).

#### 4.1.2 Reversed-contingency task

A separate cohort of 20 mice was trained on a reversed-contingency variant of the task. All training procedures, behavioral parameters, and experimental protocols were identical to those described here: (Appendix 2 of the International Brain Laboratory, 2021), with one modification: **the wheel-stimulus mapping was inverted**.

In this reversed-contingency task, wheel rotations moved the visual stimulus in the opposite direction to that of the original task. For example, a clockwise wheel rotation that would move the visual stimulus rightward in the original task instead moved it leftward in the reversed task. This manipulation reversed the contingency between motor action and visual stimulus movement, while preserving all other aspects of the original task (i.e. animal shaping protocol, reward magnitude, and trial structure).

#### 4.1.3 Electrophysiological data

Spike counts were obtained by summing spikes across time for both wiggle and non-wiggle trials. The neural data matrix had dimensions *N* × *T* × *K*, corresponding to *N* neurons, *T* time bins, and *K* trials, with indices *n* = 1, …, *N* and *t* = 1, …, *T*. The bin size was 0.02 seconds. The input to the logistic regression decoder was the single-trial neural activity, given by:

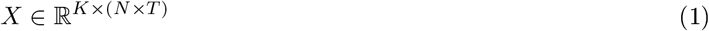

Each input vector to the logistic regression decoder is a flattened neural population activity per trial reshaped to a row vector of length *N* · *T* :

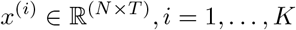

### 4.2 Quantitative and statistical analysis

#### 4.2.1 Inclusion criteria for behavioral analysis

##### Trial

Trials with a go-cue onset and visual stimulus onset within 0.05 seconds and a reaction time between 0.08 and 1.20 seconds were included in this analysis.

##### Session

Sessions with a minimum of 1 trial in each group were included in this analysis unless otherwise stated. There were 5 groups based on number of changes in wheel direction: (1) *k* = 0, (2) *k* = 1, (3) *k* = 2, (4) *k* = 3, and (5) *k* ≥ 4.

#### 4.2.2 Inclusion criteria for neural analysis

##### Insertion

Electrode insertions with a maximum of 80 microns of drift and a minimum of 10 units were included in this analysis.

#### 4.2.3 Inclusion criteria for pupil analysis

##### Session

Pupil analysis was restricted to the brain-wide map dataset (459 sessions from 139 mice), in which the video data were processed and quality-controlled: Brain-wide map quality control video spreadsheet (The International Brain Laboratory et al., 2025) Sessions were excluded if video data were missing or timestamps were corrupted. This removed 21.5% of data (99 of 459 sessions). Sessions were included based on qualitative visual inspection of pupil tracking overlaid on raw video frames. Specifically, sessions were retained when the pupil was clearly visible, image contrast and brightness were sufficient for reliable tracking, and the eyelid was not substantially closed. This removed 29.2% of data (105 of 360 sessions). The reliability of the Lightning Pose method used to generate these estimates has been validated using two post-hoc quality metrics (Biderman and Whiteway et al., 2025). First, the correlation between the vertical and horizontal pupil diameters was high (Pearson’s r = 0.88 ± 0.01, mean ± SEM). Second, the trial-to-trial consistency (defined as the variance of the mean pupil diameter trace divided by the variance of the mean-subtracted traces across all trials) was high (0.74 ± 0.08). Example frames illustrating included and excluded sessions are provided in **Figure S14**.

#### 4.2.4 Psychometric curves

To obtain a psychometric curve, data points were fitted with the following parametric error function, using a maximum likelihood procedure:

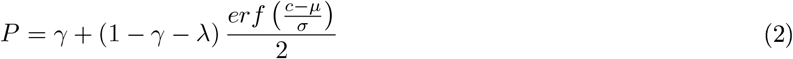

*P* probability of rightward choice

*c* visual stimulus contrast (%)

*γ* lapse rate for left visual stimuli

*λ* lapse rate for right visual stimuli

*µ* response bias

*σ* contrast threshold

A detailed psychometric curve fitting procedure is in Protocol 2 (The International Brain Laboratory, 2020b).

The **lapse rate** (*λ*) is defined as the probability of an incorrect response independent of the visual stimulus contrast (Prins et al, 2012). It captures the difference between observed performance and perfect performance at the asymptotes (Gupta et al, 2024).

#### 4.2.5 Definition of wiggles

Wiggles are defined as movements with one or more directional changes between visual stimulus onset and feedback time. A directional change is defined as a sign change in the wheel velocity (deg/sec) vs time (sec) data.

#### 4.2.6 Definition of wiggle speed

Wiggle speed is defined as the total angular distance between each pair of directional changes divided by the total wiggle duration in wheel position (deg) vs time (sec):

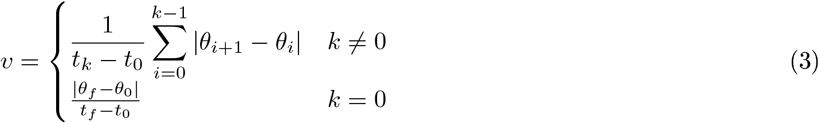

*k* number of changes in wheel direction

*θ*_0_ wheel position of motion onset (deg)

*θ*_*f*_ wheel position of feedback (deg)

*t*_0_ time of motion onset (sec)

*t*_*f*_ time of feedback (sec)

##### Quantification of temporal frequency

To connect wheel movements to visual stimulus motion, we first converted wheel speed (in wheel degrees per second) to visual stimulus speed (in visual degrees per second) using a fixed conversion factor (1 wheel degree ≈ 2 visual degrees; 0.3 wheel radians ≈ 35 visual degrees). We then divided visual stimulus speed by the spatial frequency of the Gabor patch (0.1 cycles per degree) to obtain the effective temporal frequency experienced by the mouse. This temporal frequency reflects how rapidly the stimulus pattern drifted across the retina as the mouse wiggled the wheel. Thus, visual temporal frequency is defined as wiggle speed divided by stimulus spatial frequency.

#### 4.2.7 Definition of motion energy

Motion energy is defined as the time-averaged speed squared across each pair of directional changes in wheel position (deg) vs time (sec):

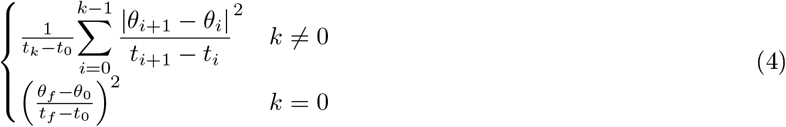

Motion energy is in units of deg^2^/sec^2^ and ranges from 500 to 80,500 deg^2^/sec^2^. For each trial, we summed the squared positional changes between consecutive directional changes divided by their corresponding time intervals (deg^2^/sec). This value was then normalized by the total wiggle duration (sec) to obtain a time-averaged measure of movement intensity. To exclude ballistic movements that often follow wiggles in *k* ≠ 0 trials, we performed analysis up to *k* ≠ 1 intervals rather than *k* intervals.

### 4.2.8 Definition of visual angle

Visual angle is the perceived size of a stimulus given by:

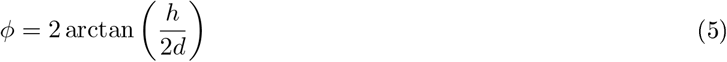

*h* height of monitor (cm)

*d* distance between mouse and monitor (cm)

*φ* visual angle (deg)

Mice viewed a Gabor patch on a screen size of 246 mm diagonal active display area and were positioned 8 cm away from the screen to achieve 102 visual degrees azimuth.^1^ The Gabor patch size is 19.7 cm if *φ* = 102° and *d* = 8 cm.

Visual angle represents the size of an object’s image on the retina. To obtain visual angle, we converted from wheel degrees to visual degrees. 35 visual degrees equals 17.2 wheel degrees (0.3 radians). Thus, 2 visual degrees correspond to about 1 wheel degree.

#### 4.2.9 Weighted linear regression analysis

First, we binned the wiggle speed data across sessions per mouse to obtain a distribution of the proportion correct for a given visual contrast level. Each data point represented one session. We performed a logistic regression between proportion correct and wiggle speed across 8 octiles per mouse. Second, we pooled the median proportion correct per wiggle speed bin across 213 mice. Each data point represented one mouse. We performed a logistic regression between median proportion correct and motion energy across 8 octiles in 213 mice.

#### 4.2.10 Pupil data analysis

To examine whether wiggle behavior could be explained by arousal, we performed four complementary statistical analyses using pupil diameter as a proxy for arousal state.

##### Relationship between wiggle metrics and pupil diameter

We assessed the relationship between wiggle metrics and pupil diameter at the session level. For each session, we calculated the median value of each wiggle metric and the median pupil diameter across all wiggle trials at 6.25% contrast. Wiggle speed data were binned into octiles to be consistent with the behavioral analysis in **Figure 3a** and pupil-controlled analysis in **Figure 5e**. We then performed weighted linear regressions between the median value of each wiggle metric (speed, amplitude, number of direction changes, or duration) and the median pupil diameter across 255 sessions from 93 mice, where sessions were weighted by the number of wiggle trials at 6.25% contrast.

##### Pupil-controlled analyses

We performed a pupil-controlled analysis using two approaches. In one approach, we binned trials into narrow pupil diameter bins (0.05 a.u. from 0.10 to 0.45). Within each pupil bin, we partitioned trials into octiles of wiggle speed. We then performed weighted linear regression between wiggle speed octile and proportion correct. In the second approach, we used a linear mixed-effects model with wiggle speed and pupil diameter as fixed effects and mouse identity as a random intercept. The outcome variable was proportion correct at the octile level (each session contributed eight observations, one per wiggle speed octile).

##### Classification of session types

We classified sessions as benefiting, neutral, or non-benefiting based on the within-session relationship between wiggle behavior and low-contrast accuracy. For each session, we grouped trials at 6.25% contrast by number of wheel direction changes (binned as 0, 1, 2, 3, or ≥4) and calculated the proportion correct for each bin. We then performed weighted linear regression between number of direction changes and proportion correct, where weights reflected the number of trials per bin. Sessions with a positive Pearson correlation (r > 0.3) between wiggling and accuracy were classified as benefiting, those with a negative correlation (r < −0.3) were classified as non-benefiting, and those with weak correlations (−0.3 ≤ r ≤ 0.3) were classified as neutral.

##### Comparison of pupil diameter distributions across session types

We compared pupil diameter distributions across session types (benefiting, neutral, non-benefiting) using Kruskal-Wallis tests and one-way ANOVA.

##### Relationship between wiggle speed and pupil diameter by session type

We examined whether the relationship between wiggle speed and pupil diameter differed across session types. We binned trials into wiggle speed octiles within each session and calculated the median wiggle speed and the median pupil diameter for each octile. We then used mixed-effects linear regression to model median pupil diameter as a function of median wiggle speed, with session as a random effect. Each observation in the model represented one octile within one session, so each session contributed eight observations. Separate models were fit for benefiting, neutral, and non-benefiting sessions.

#### 4.2.11 Computation of Temporal Contrast Sensitivity

A behavioral d’ was used to measure temporal contrast sensitivity (TCS):

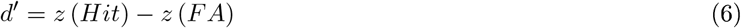

*z*(*Hit*) z-score of the probability of hits

*z*(*FA*) z-score of the probability of false alarms

Temporal contrast sensitivity is a function of temporal frequency, spatial frequency, and background mean illumination (screen brightness). Screen brightness was not well-standardized across laboratories and no published value for illumination is listed.^1^ On average, we determined that the screen brightness was 29.4 cd m^− 2^. This means that log (29.4) = 1.47 log cd m^− 2^. Therefore, we used the data under photopic (1.80 log cd m^-2^) conditions (See **Figure 2b** of Umino et al, 2008; **Figure 6f** of Umino et al, 2018) for details of mouse temporal contrast sensitivity analysis).

## Data Availability

Please follow these instructions to download the data. There are visualizations available to understand the data (Birman et al, 2024).

## Code Availability

Please visit our Github repository to access the code used to produce the main results and figures presented in this manuscript. This International Brain Lab Github repository also contains code for brain-wide neural data analysis.

## Acknowledgements

This research was supported by the Gatsby Charitable Foundation (GAT3755) and in part by grant NSF PHY-2309135 to the Kavli Institute for Theoretical Physics (KITP). Computational resources were provided by University College London. We thank Alec Ortiz and Sylvia Schröder for perturbation experiment data. We thank Thomas Mrsic-Flogel, Peter Latham, Samuel Solomon, Adil Khan, Dimitar Kostadinov, and David Kleinfeld for comments on the behavioral and neural analysis. We thank Matthew Whiteway for comments on the pupil analysis. We thank Joel E. Cohen for comments on the statistical analysis. We thank Andrew Peters, Viktor Roytman, and Alvin Zhang for comments on the manuscript. Most of all, we are indebted to Peter Dayan and Laurence F. Abbott for seeing this manuscript to completion.

## Author Contributions

N.G. conceived the project, performed the data analysis, and wrote the manuscript. A.Y.Y. trained mice on the reversed contingency task and collected the corresponding perturbation data.

## Declaration of generative AI and AI-assisted technologies in the writing process

During the preparation of this work, we used Claude and ChatGPT to revise text and debug code. After using this tool/service, we reviewed and edited the content as needed and we take full responsibility for the content of the publication.

## Notes

https://www.internationalbrainlab.com/data

### Competing Interest Statement

The authors have declared no competing interest.

### Summary of Updates

Pupil analysis revisions; Supplemental files updated.

