## Supplement for "Active sensing during a visual perceptual decision-making task"

30 March 2026

[1] Sainsbury Wellcome Centre for Neural Circuits and Behaviour, University College London, London, UK
[2] The International Brain Laboratory

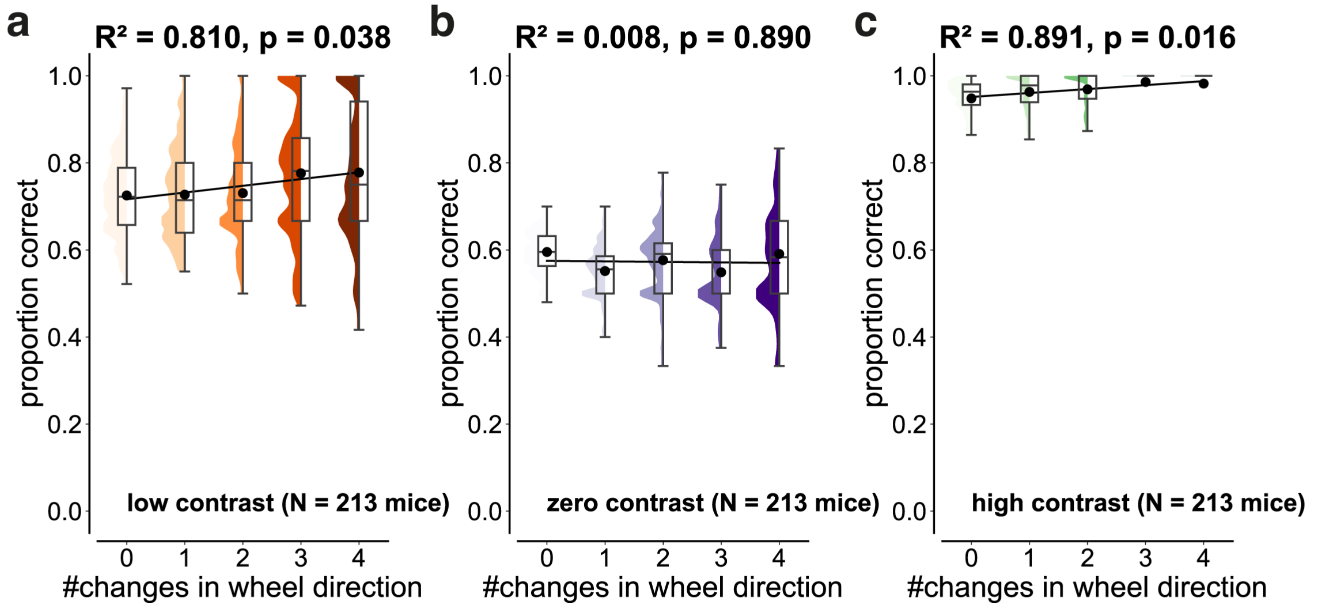


Figure S1: **The relationship between wiggle behavior and accuracy is a function of visual contrast.**

**(a-c)** Distributions of proportion correct as a function of number of changes in wheel direction ($k$) for a given visual contrast. Data were pooled across 213 mice and only sessions with $\geq 1$ trial per bin were included. Titles denote the $R^2$ and p-value of a weighted linear regression analysis between the median change in proportion correct and $k$.

**(a)** Accuracy increased with $k$ in 6.25% contrast trials (linear regression; $R^2 = 0.810$, $p = 0.038$, Cohen's $d = 0.56$, $N = 213$ mice).

**(b)** Accuracy was not significantly dependent on $k$ in 0% contrast trials (linear regression; $R^2 = 0.008$, $p = 0.890$, Cohen's $d = 0.30$, $N = 213$ mice).

**(c)** Accuracy increased with $k$ in 100% contrast trials (linear regression; $R^2 = 0.891$, $p = 0.016$, Cohen's $d = 0.93$, $N = 213$ mice).

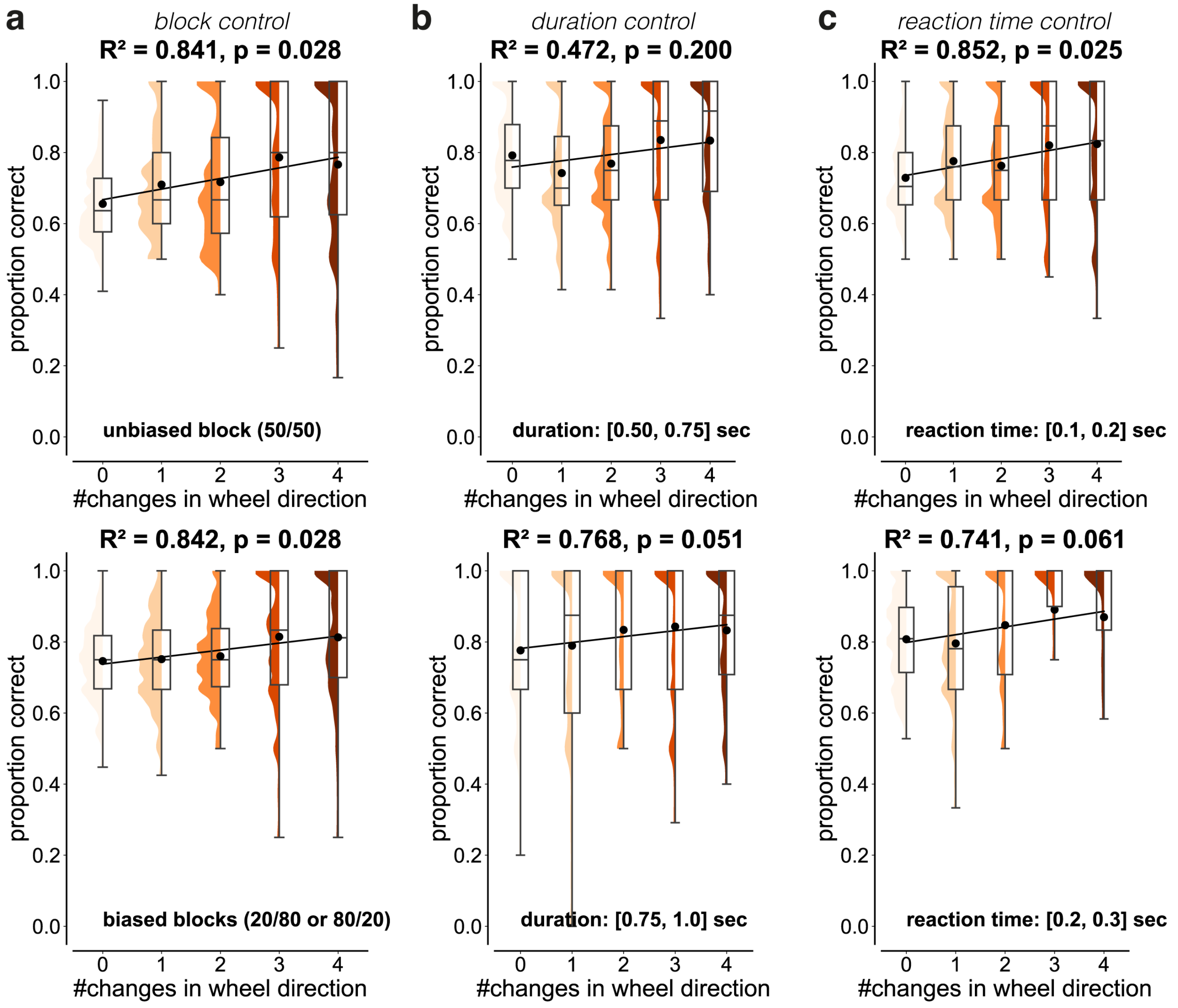


Figure S2: **Wiggle behavior is not explained by differences in block, trial duration, and reaction time.** We performed a weighted linear regression analysis across 213 mice during 6.25% contrast trials as described in **Figure S1**.

**(a) Block control:** Accuracy increased with $k$ in unbiased and biased blocks (**unbiased:** $R^2 = 0.841$, $p = 0.028$, Cohen's $d = 0.66$; **biased:** $R^2 = 0.842$, $p = 0.028$, Cohen's $d = 0.66$).

**(b) Duration control:** Accuracy increased with $k$ in trials of similar durations (**0.50 − 0.75 sec:** $R^2 = 0.472$, $p = 0.200$, Cohen's $d = 0.45$; **0.75 − 1.0 sec:** $R^2 = 0.768$, $p = 0.051$, Cohen's $d = 0.32$), although these effects did not reach statistical significance.

**(c) Reaction time control:** Accuracy increased with $k$ in trials of similar reaction times (**0.1 − 0.2 sec:** $R^2 = 0.852$, $p = 0.025$, Cohen's $d = 0.60$; **0.2 − 0.3 sec:** $R^2 = 0.741$, $p = 0.061$, Cohen's $d = 0.31$).

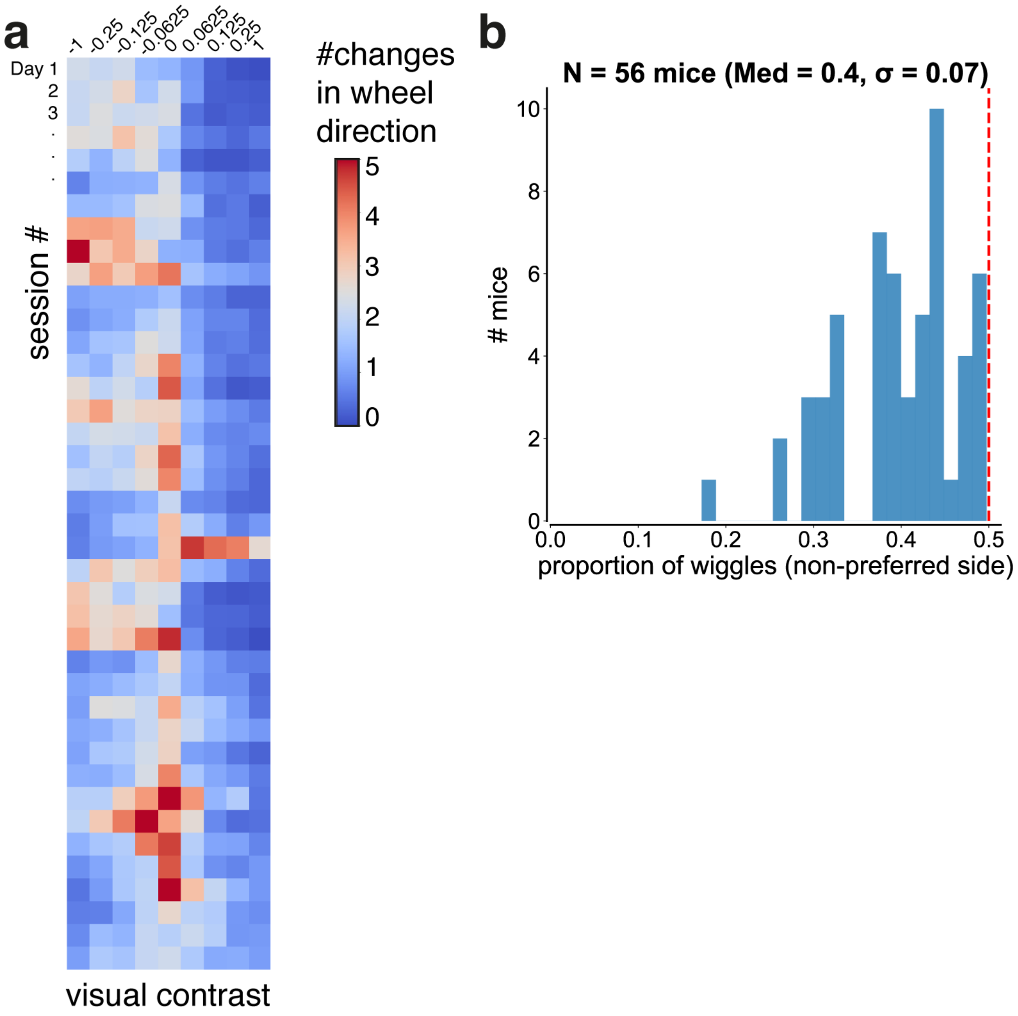


Figure S3: **Mice with asymmetric wiggle behavior preferentially wiggle on the higher-accuracy side.**

**(a)** Heat map showing the mean number of changes in wheel direction ($k$) as a function of visual contrast and training day for a representative asymmetric good wiggler (NYU-04).

**(b)** Distribution of the fraction of wiggle trials directed to the non-preferred side across mice with asymmetric wiggle behavior (N = 56 mice). The non-preferred side was defined as the side associated with lower accuracy during 6.25% contrast trials. The median fraction of non-preferred wiggles was 0.40 (IQR: 0.36 – 0.44), significantly below chance (0.5; one-sample Wilcoxon signed-rank test, $p < 0.001$). The red dashed line denotes chance level (0.5).

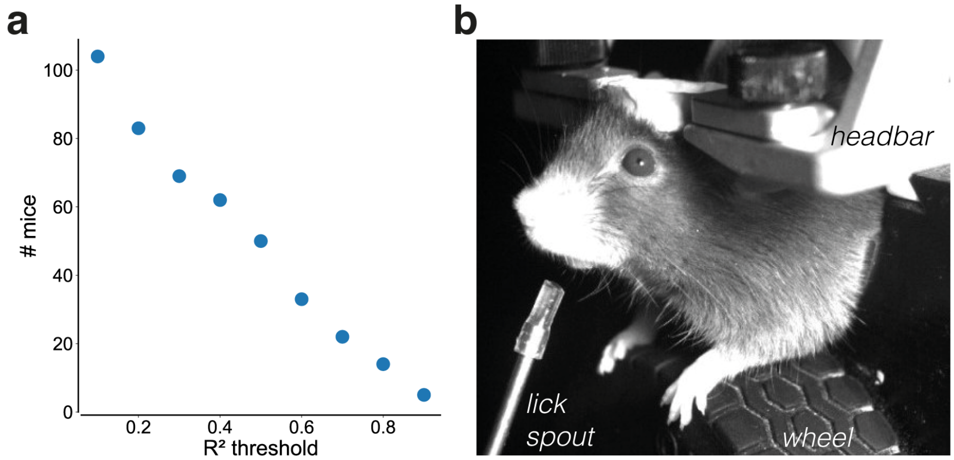


Figure S4: **The number of mice benefiting from wiggle behavior depends on the $R^2$ threshold.**

**(a)** Number of mice with a positive relationship between performance and wiggle behavior as a function of goodness-of-fit. Scatter plot showing the number of mice with a positive regression slope between proportion correct and the number of directional changes ($k$) during 6.25% contrast trials, plotted against the $R^2$ threshold.

**(b)** Example video frame illustrating a mouse using the steering wheel. Photo credit: Petrina Lau.

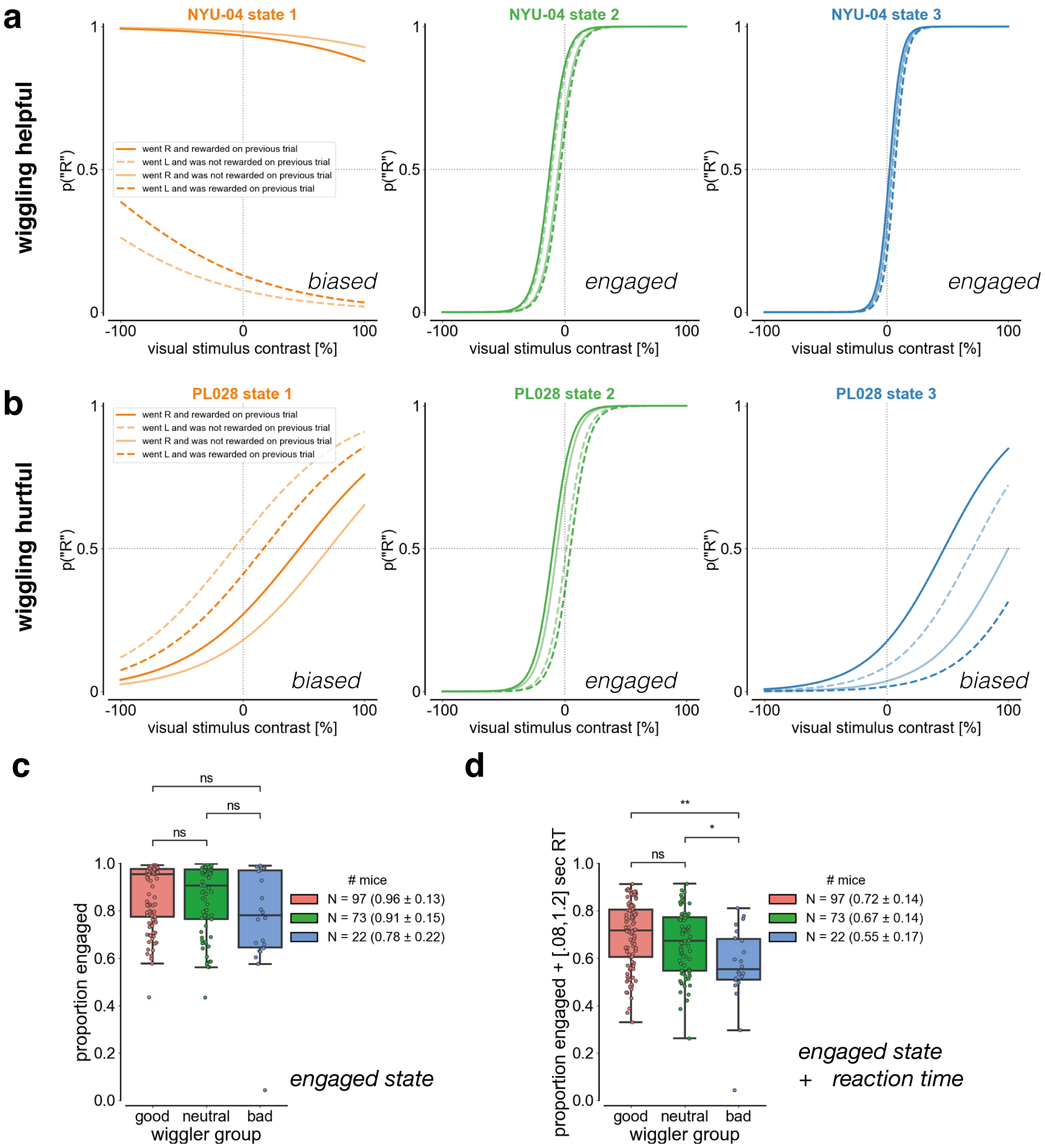


Figure S5: **Reaction time differentiates mice for whom wiggle behavior is beneficial versus detrimental to accuracy.** Wheel movements were classified as engaged, biased, or disengaged using a three-state GLM-HMM fit separately for each mouse (Ashwood et al, 2022). Mice were further grouped based on the relationship between proportion correct and wiggle behavior: **good wigglers** (positive slope), **neutral wigglers** (near-zero slope), and **bad wigglers** (negative slope).

**(a)** Example of three GLM-HMM states for a representative good wiggler (NYU-04).

**(b)** Example of three GLM-HMM states for a representative bad wiggler (PL028).

**(c)** Proportion of wheel movements classified as engaged as a function of wiggler group. This proportion did not differ significantly between good and bad wigglers (Mann-Whitney U test with Bonferroni correction; $\alpha = 0.016$, U = 1,325, p = 0.078, 95% CI: [11, 25]).

**(d)** Proportion of wheel movements classified as engaged with reaction times between 0.08 and 1.2 seconds as a function of wiggler group. This proportion was significantly higher in good wigglers (Mann-Whitney U test with Bonferroni correction, $\alpha = 0.0167$, U = 1,548, p = 0.001, 95% CI: [10, 24], corrected Cohen's d = 0.87).

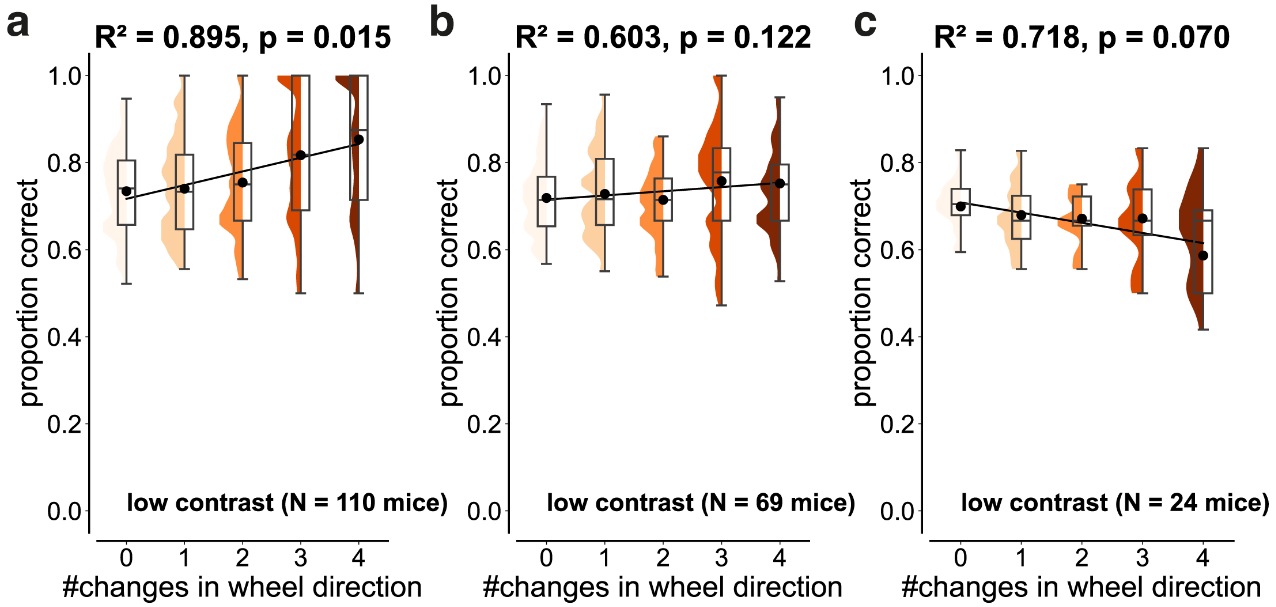


Figure S6: **The effect of wiggle behavior is heterogeneous across mice.** We performed a weighted linear regression analysis across subsets of mice during 6.25% contrast trials as described in **Figure S1.**

**(a) Good wigglers:** There are 110 mice with a monotonically increasing relationship between $k$ and low-contrast accuracy ($R^2 = 0.895$, p = 0.015, Cohen's d = 1.09).

**(b) Neutral wigglers:** There are 69 mice with a negligible relationship between $k$ and low-contrast accuracy ($R^2 = 0.603$, p = 0.122, Cohen's d = 0.90).

**(c) Bad wigglers:** There are 24 mice with a monotonically decreasing relationship between $k$ and low-contrast accuracy ($R^2 = 0.718$, p = 0.070, Cohen's d = 0.86).

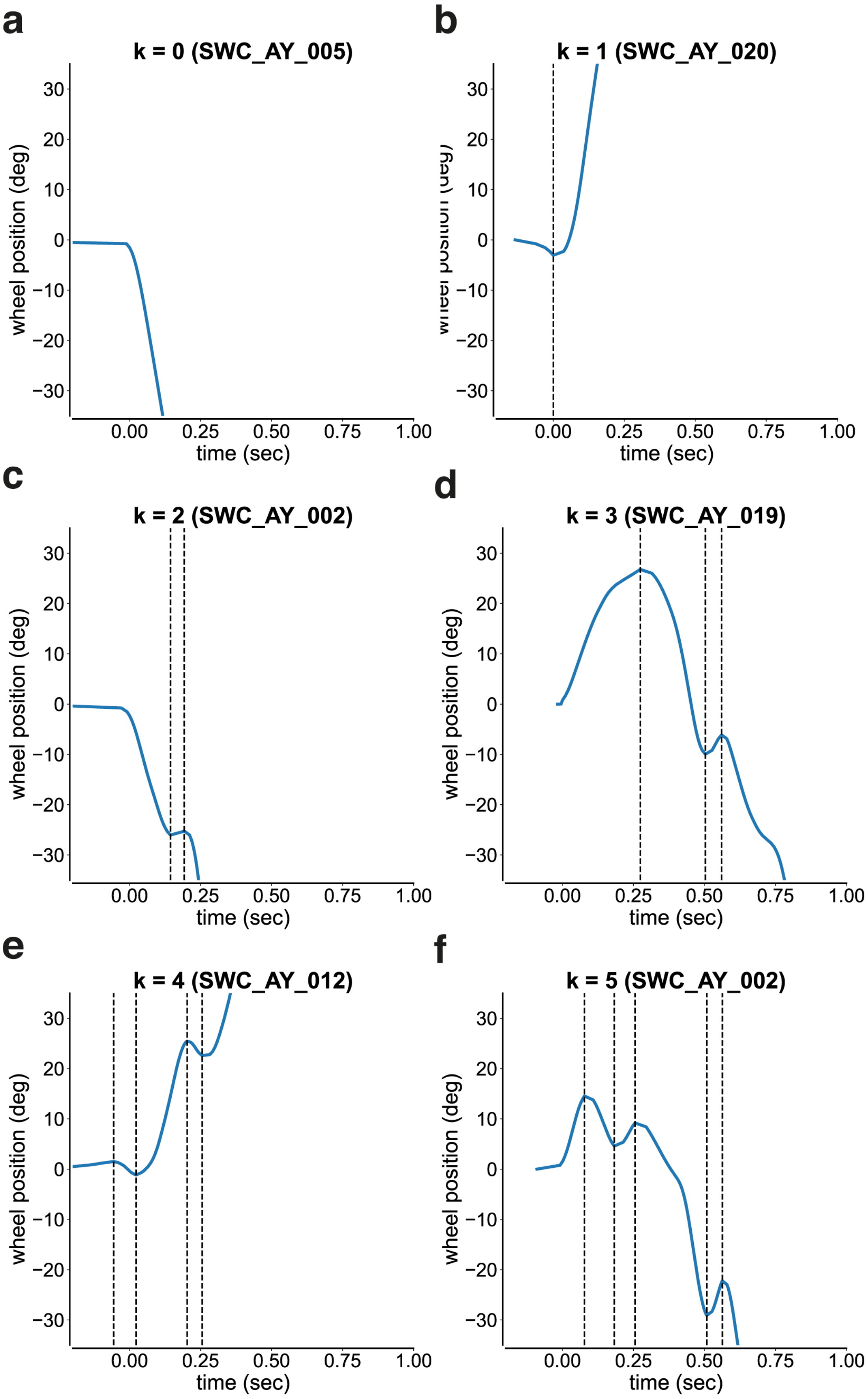


Figure S7: **Wheel movements are comparable between the reversed and standard contingency tasks.**

**(a-f)** Single trial examples of wheel position (deg) over time (sec) for 6.25% contrast trials in the reversed-contingency control task, shown for trials with **(a)** $k = 0$, **(b)** $k = 1$, **(c)** $k = 2$, **(d)** $k = 3$, **(e)** $k = 4$, and **(f)** $k = 5$ directional changes. Trials are aligned to motion onset, as in **Figure 2**.

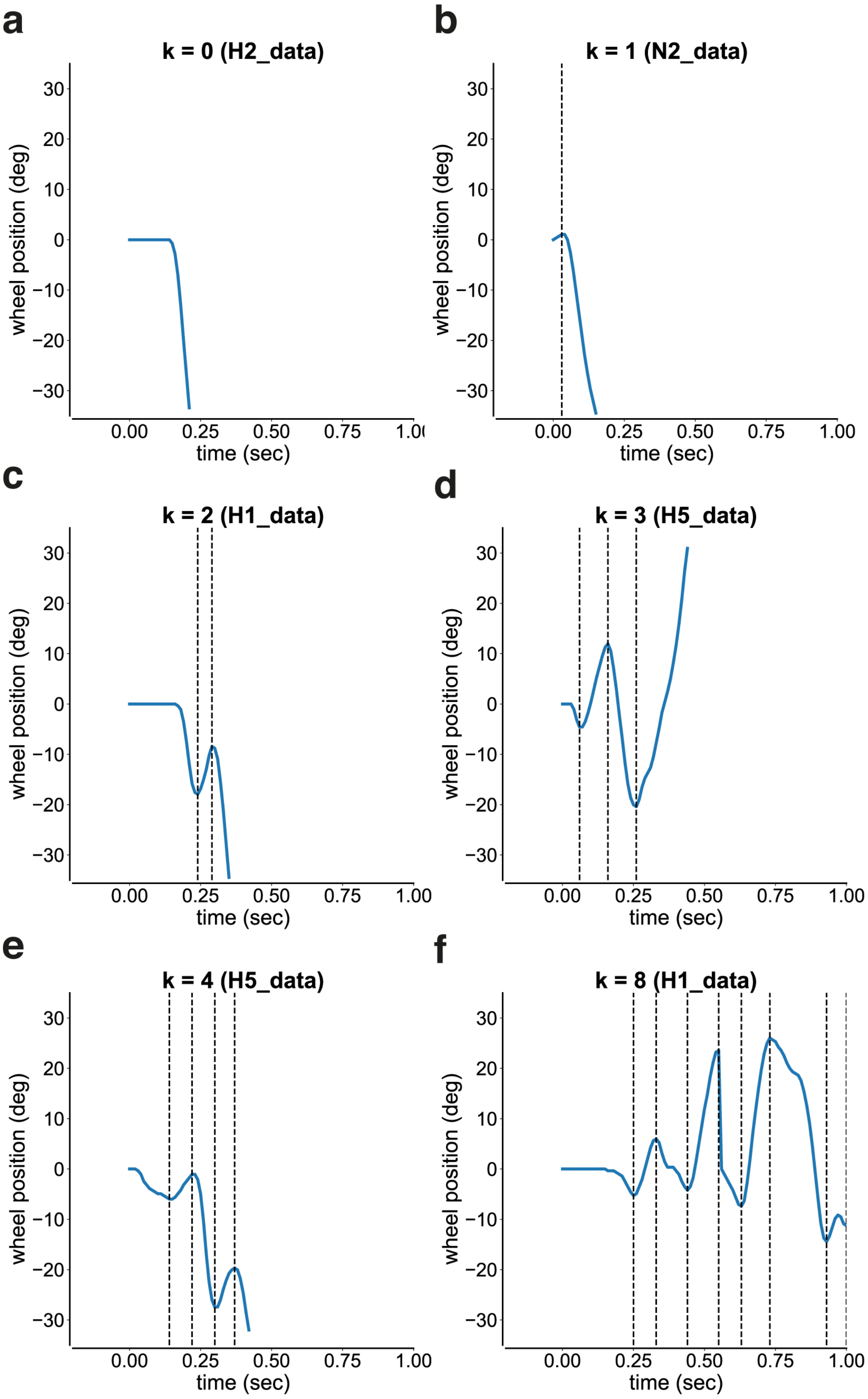


Figure S8: **Wheel movements are comparable across spatial frequency conditions.**

**(a-f)** Single trial examples of wheel position (deg) over time (sec) for 4% contrast trials in the spatial frequency control, shown for trials with **(a)** $k = 0$, **(b)** $k = 1$, **(c)** $k = 2$, **(d)** $k = 3$, **(e)** $k = 4$, and **(f)** $k = 8$ directional changes. Trials are aligned to visual stimulus onset.

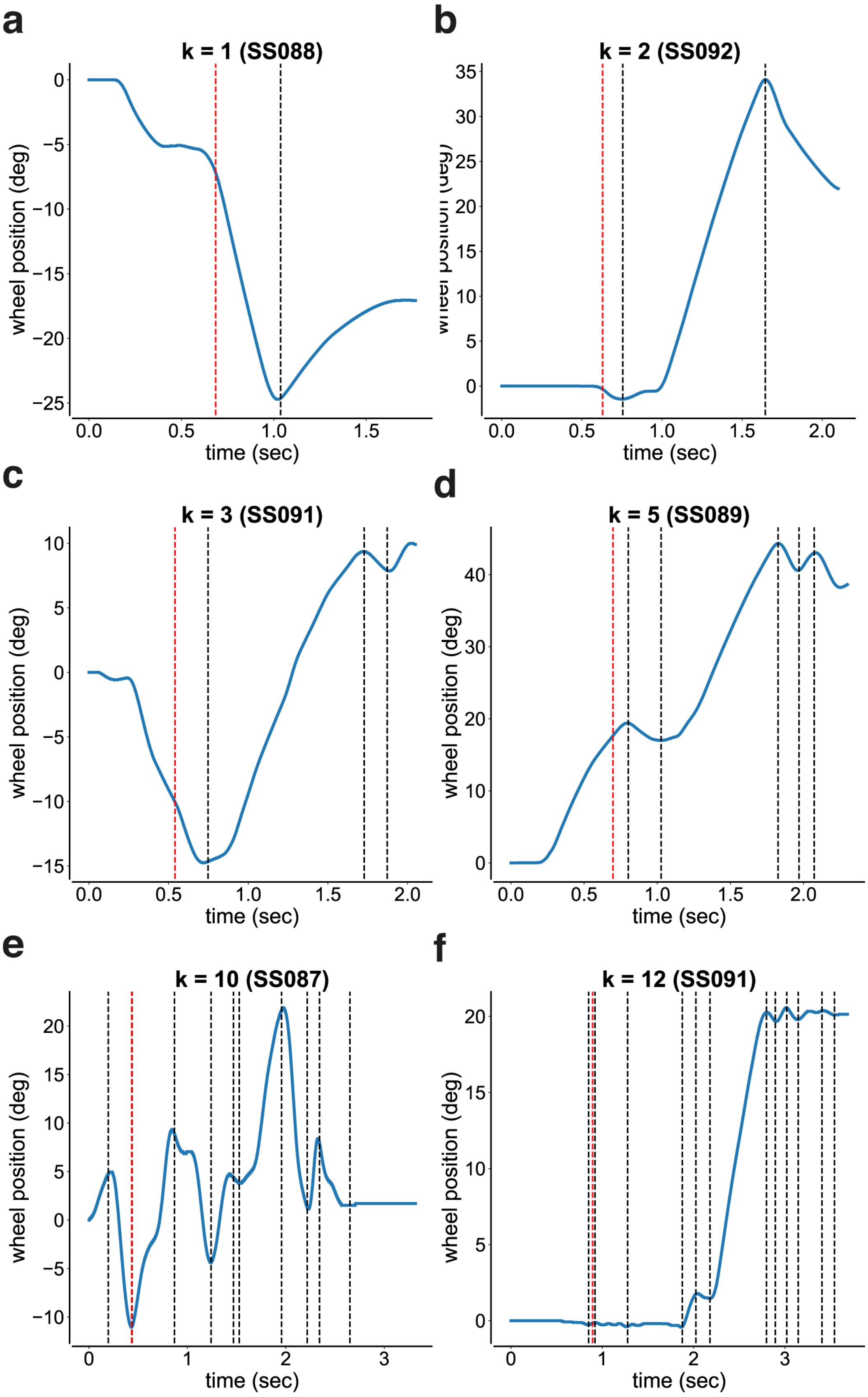


Figure S9: **Wheel movements are comparable in open-loop and closed-loop tasks.**

**(a-f)** Single trial examples of wheel position (deg) over time (sec) for 25% contrast trials in the open-loop and closed-loop tasks, shown for trials with **(a)** $k = 1$, **(b)** $k = 2$, **(c)** $k = 3$, **(d)** $k = 5$, **(e)** $k = 10$, and **(f)** $k = 12$ directional changes. Trials are aligned to visual stimulus onset. The red dashed line indicates go-cue onset, which is when the wheel became coupled to the visual stimulus.

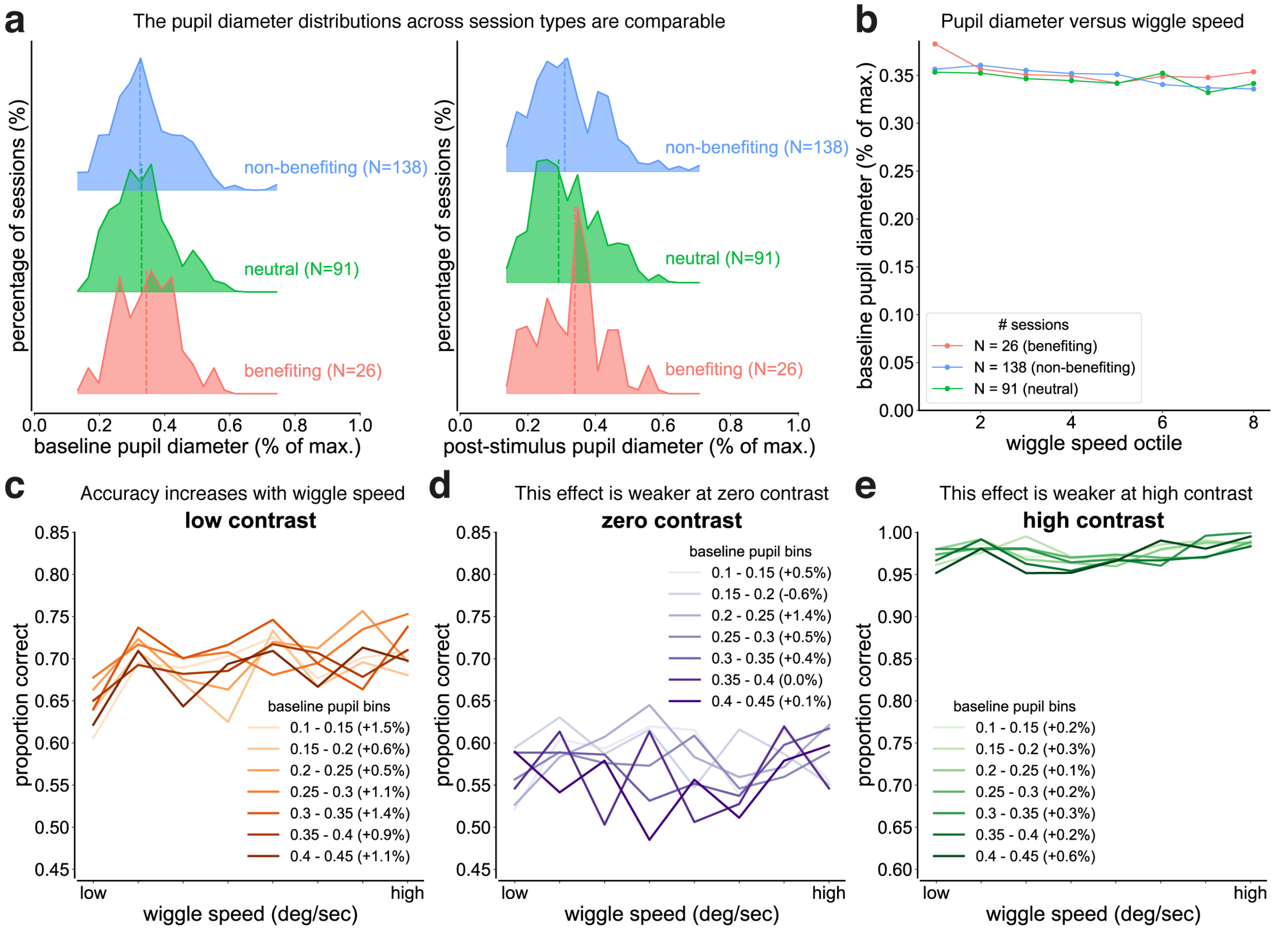


Figure S10: **Wiggle behavior enhances low-contrast accuracy independently of arousal (additional analyses using baseline pupil diameter).** Analysis performed across 255 sessions from 93 mice as described in **Figure 5.**

**(a)** Distributions of normalized baseline pupil diameter during 6.25% contrast trials by session type, shown as vertically offset (ridgeline) histograms, with each distribution normalized to percentage within condition. Sessions were classified based on the within-session relationship between wiggle behavior (number of wheel direction changes) and low-contrast accuracy: benefiting ($r > 0.3$, $N = 26$), neutral ($-0.3 \leq r \leq 0.3$, $N = 91$), or non-benefiting ($r < -0.3$, $N = 138$). The pupil diameter distributions are comparable across session types (Kruskal-Wallis test, $p = 0.84$). The dashed lines represent medians.

**(b)** Relationship between wiggle speed and baseline normalized pupil diameter by session type (approach identical to **Figure 5h**). Trials were binned into wiggle speed octiles within each session. Mixed-effects linear regression with session as a random effect showed no significant relationship in benefiting sessions ($\beta = -0.002$, $p = 0.20$), but a small significant decrease in neutral ($\beta = -0.005$, $p = 0.02$) and non-benefiting sessions ($\beta = -0.003$, $p < 0.001$).

**(c-e)** Relationship between wiggle speed and accuracy at matched arousal levels using baseline pupil diameter (approach identical to **Figures 5e–g**). Baseline normalized pupil diameter was binned in 0.05 a.u. increments. Within each pupil bin, trials were partitioned into octiles of wiggle speed, and accuracy was computed for each octile.

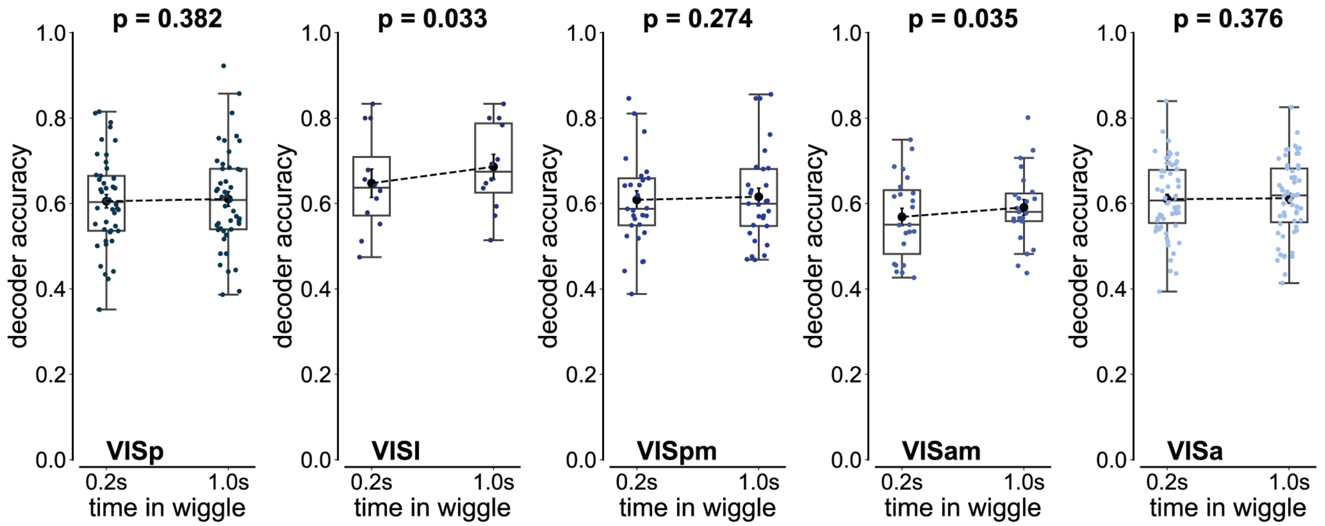


Figure S11: **The effect of wiggle duration on stimulus side decoding accuracy varies across visual cortical regions.** Brain region abbreviations and colors follow the Allen Institute Brain Atlas (VISp: primary visual area; VISl: lateral visual area; VISpm: posteromedial visual area; VISam: anteromedial visual area; VISa: anterior visual area). All statistical comparisons were performed using Wilcoxon signed-rank tests ($\alpha = 0.05$). Some higher-order visual areas showed significant improvements in decoding accuracy for long versus short wiggle trials: VISl ($p = 0.033$, $N = 12$ sessions, median improvement: 3.8%) and VISam ($p = 0.035$, $N = 23$ sessions, median improvement: 2.2%). In contrast, primary visual cortex showed no significant change in decoding accuracy: VISp ($p = 0.382$, $N = 45$ sessions, median improvement: -0.7%). Similar null effects were observed in other higher-order visual areas: VISpm ($p = 0.274$, $N = 29$ sessions, median improvement: 0.7%) and VISa ($p = 0.376$, $N = 98$ sessions, median improvement: 0.2%)

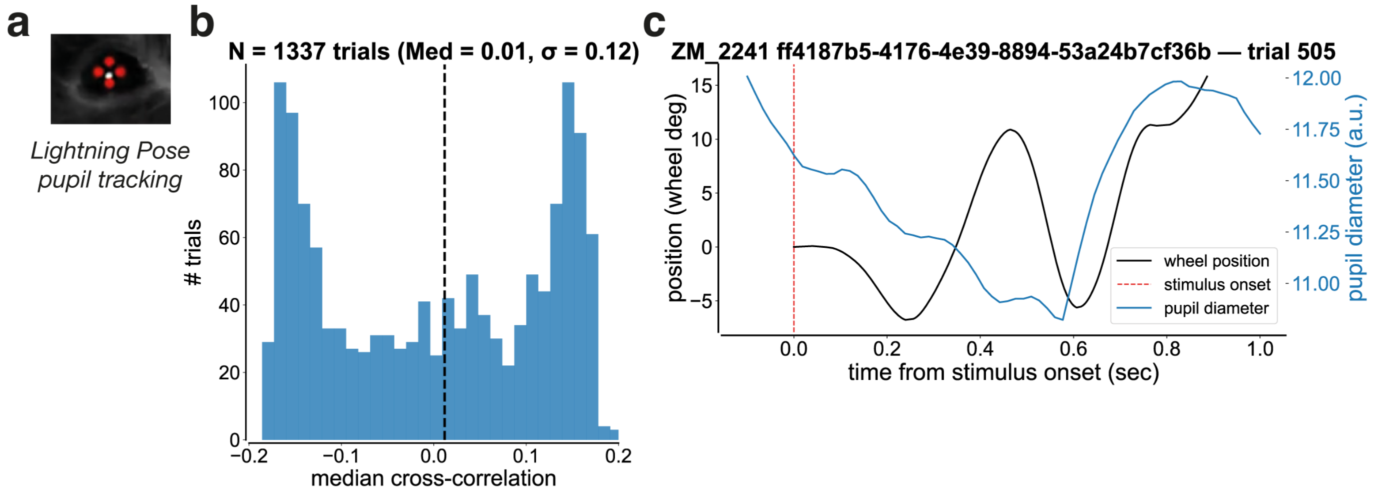


Figure S12: **Wiggle behavior is not correlated with arousal.**

**(a)** Example video frame illustrating pupil tracking. Pupil diameter was defined as the vertical distance between the top and bottom pupil coordinates, estimated using the Lightning Pose algorithm. Photo credit: Matthew Whiteway.

**(b)** Distribution of median cross-correlation coefficients between pupil diameter and wheel position across wiggle trials (median: 0.01, standard deviation: 0.12, range: [-0.18, 0.20], N = 1,337 wiggle trials). Cross-correlations were computed within a 500-msec window following visual stimulus onset. Negative values indicate that the pupil diameter trace lagged the wheel position trace. The black dashed line denotes the median.

**(c)** Single-trial example showing pupil diameter (blue) and wheel position (black) traces aligned to visual stimulus onset (red dashed line), illustrating the method used to compute the cross-correlations shown in **(b)**.

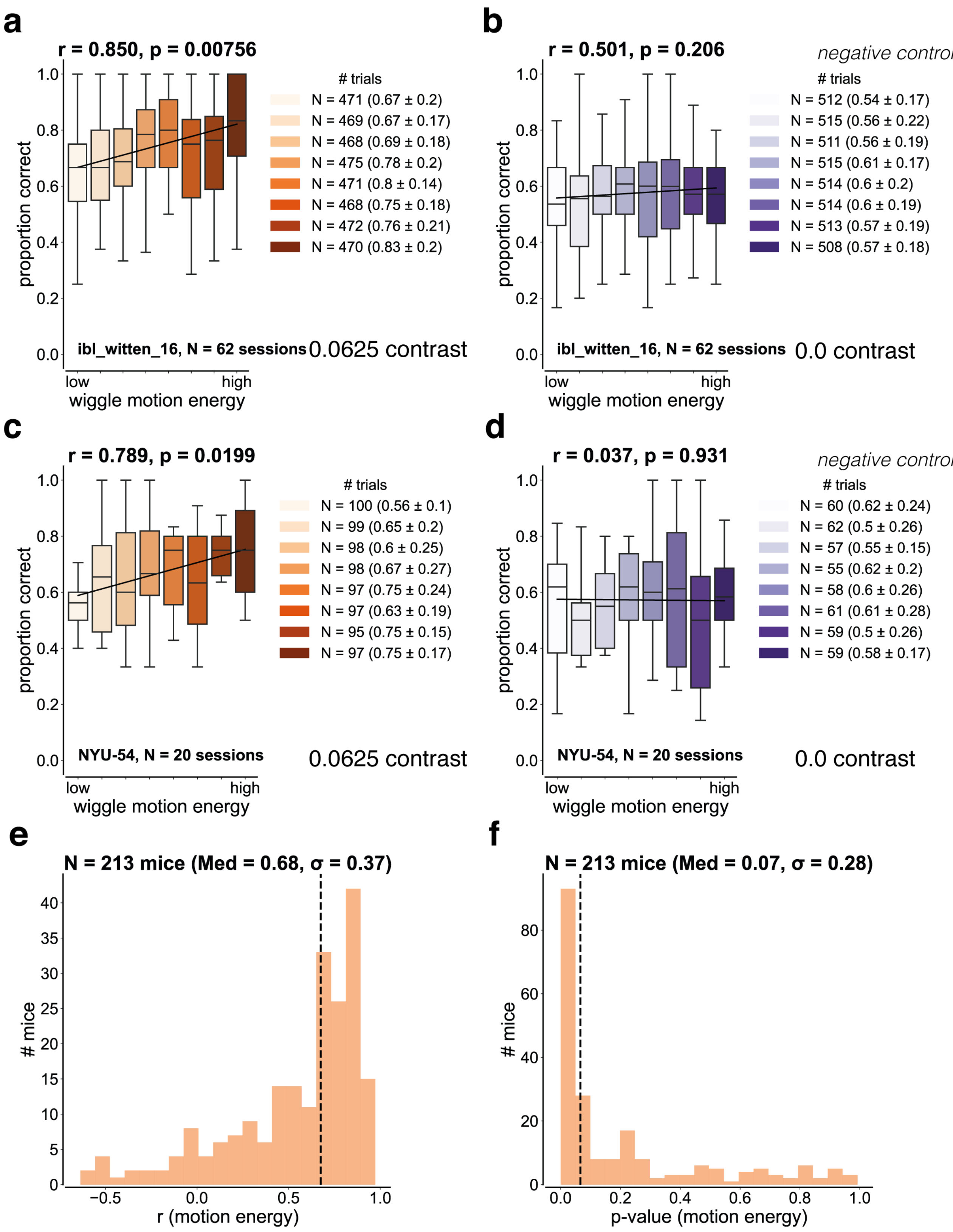


**Figure S13: A subset of mice benefits from wiggle behavior.**

**(a-d)** Distributions of proportion correct as a function of motion energy for a given contrast per mouse. Motion energy was defined as the time-averaged squared speed between successive directional changes divided by the wiggle duration, which was defined as the time between the first and last extremum. Data were pooled across sessions per mouse and binned into equal octiles; only sessions with $\geq 3$ trials per bin were included. Titles denote Pearson correlation coefficient and p-value across octile-medians.

**(a)** Accuracy increased with motion energy in 6.25% contrast trials (logistic regression; $\beta = 0.092 \pm 0.016$ SE, p = $1.11 \times 10^{-8}$, Cohen's d = 0.091; Pearson correlation of octile-medians, r = 0.850, p = 0.007; N = 62 sessions, 1 mouse: ibl_witten_16).

**(b)** Accuracy was not significantly dependent on motion energy in 0% contrast trials (logistic regression; $\beta = 0.024 \pm 0.014$ SE, p = 0.079, Cohen's d = 0.024; Pearson correlation of octile-medians, r = 0.501, p = 0.206; N = 62 sessions, 1 mouse: ibl_witten_16).

**(c)** Accuracy increased with motion energy in 6.25% contrast trials (logistic regression; $\beta = 0.122 \pm 0.033$, p = 0.00025, Cohen's d = 0.122; Pearson correlation of octile-medians, r = 0.789, p = 0.020, N = 20 sessions, 1 mouse: NYU-54).

**(d)** Accuracy was not significantly dependent on motion energy in 0% contrast trials (logistic regression; $\beta = -0.014 \pm 0.039$ SE, p = 0.72, Cohen's d = -0.014; Pearson correlation of octile-medians, r = 0.037, p = 0.931, N = 20 sessions, 1 mouse: NYU-54).

**(e)** Distribution of per-mouse Pearson correlation coefficients (r) between motion energy and proportion correct during 6.25% contrast trials (median: 0.68, standard deviation: 0.37). The black dashed line denotes the median. The distribution is significantly skewed towards positive r values (skewness: -6.17, p = $6.58 \times 10^{-10}$, N = 213 mice).

**(f)** Distribution of corresponding per-mouse p-values assessing the relationship between motion energy and proportion correct during 6.25% contrast trials (median: 0.07, standard deviation: 0.28). The black dashed line denotes the median.

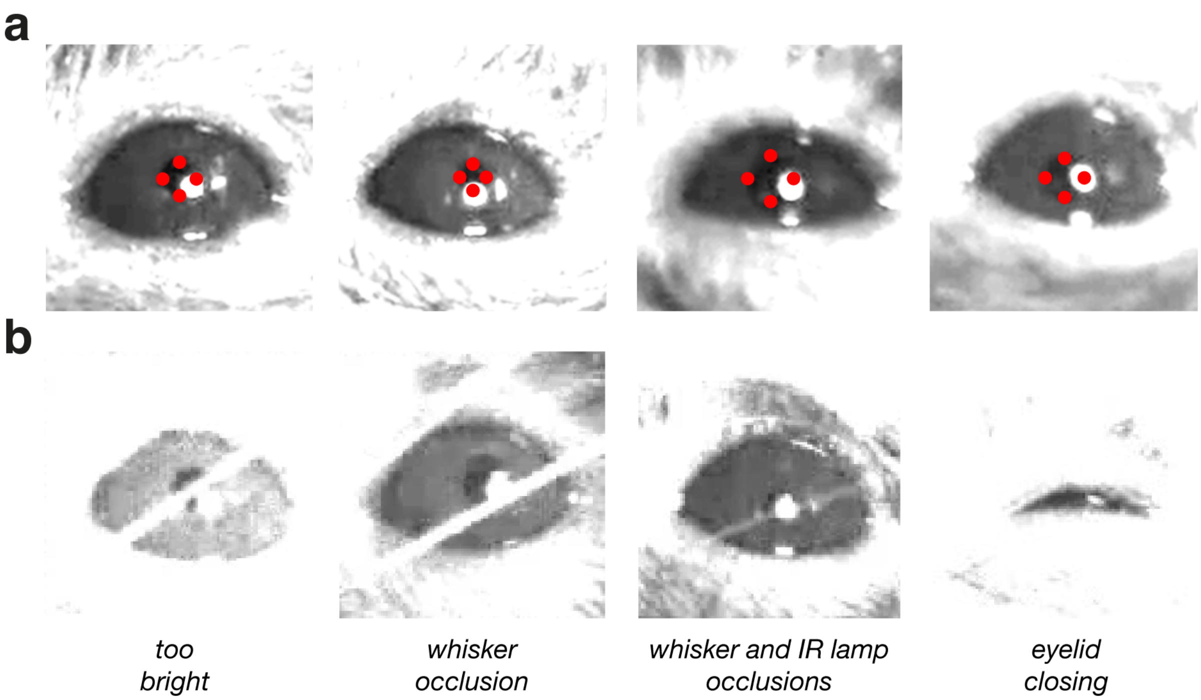


Figure S14: **The quality of pupil tracking varies across sessions.** Pupil boundary markers overlaid on raw video frames indicate pupil tracking by Lightning Pose (Biderman and Whiteway et al., 2025).

**(a)** Example video frames from included sessions, in which the pupil is clearly visible and image contrast is sufficient for reliable tracking.

**(b)** Example video frames from excluded sessions, illustrating common failure modes: poor image brightness, whisker occlusions, or a substantially closed eyelid.

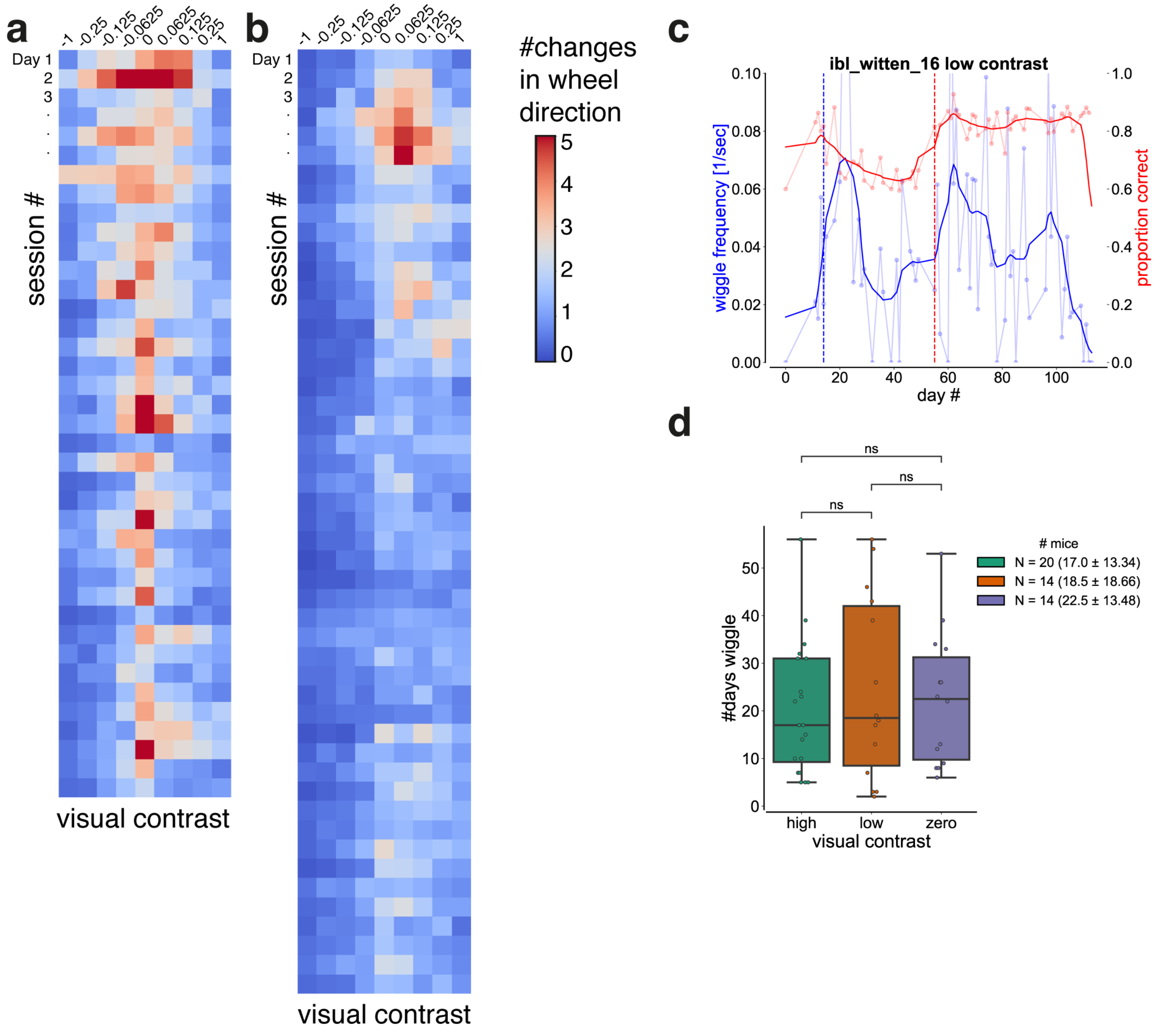


Figure S15: **Wiggle behavior emerges over training.**

**(a)** Heat map showing the mean number of changes in wheel direction ($k$) as a function of visual contrast and training day for a representative good wiggler (ibl_witten_16).

**(b)** Corresponding heat map for a representative bad wiggler (PL028).

**(c)** Proportion of trials with $k \geq 4$ (blue) plotted across training days alongside proportion correct (red) during 6.25% contrast trials for a representative good wiggler (NYU-04).

**(d)** Proportion of wiggle days as a function of visual contrast across mice. Wiggle days were defined as sessions containing at least 2% $k \geq 4$ trials, provided this criterion was met on at least two days. Only mice with a minimum of 12 sessions containing $k \geq 4$ trials were included in this analysis.

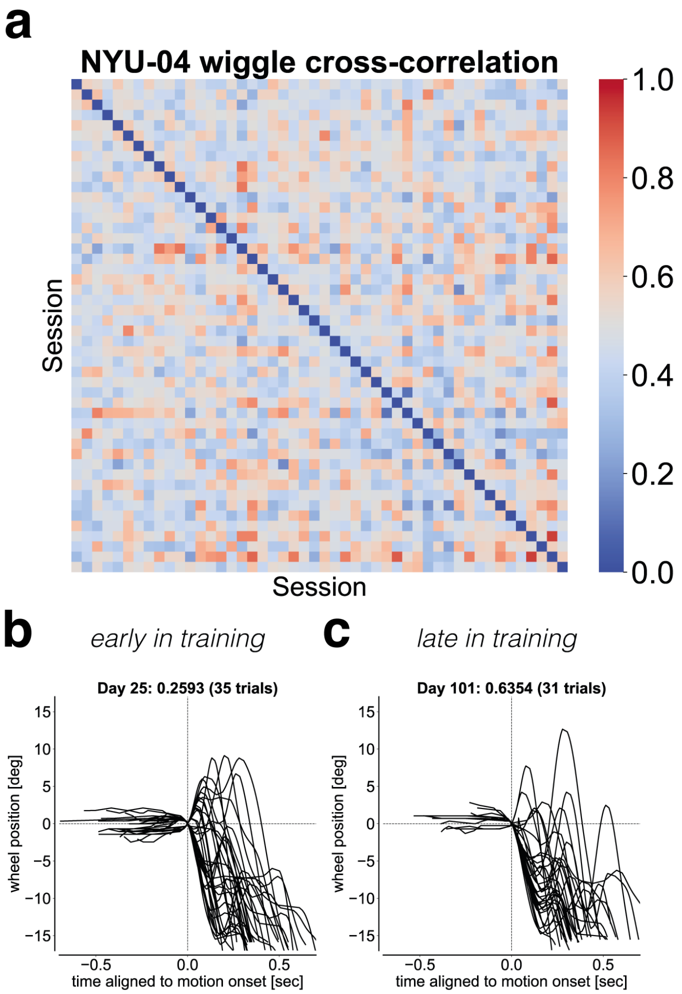


Figure S16: **Wiggle behavior is not stereotyped.**

**(a)** Cross-correlation matrix of the proportion of wiggle trials across sessions for a representative good wiggler (NYU-04). Cross-correlation coefficients were near zero, indicating no evidence for stereotyped behavior across sessions.

**(b)** Example of wiggle trials from early training (Day 25).

**(c)** Corresponding example wiggle trials from late training (Day 101).
